# Background compensation revisited: Conserved phase response curves in frequency controlled homeostats with coherent feedback

**DOI:** 10.1101/2024.06.06.597853

**Authors:** Peter Ruoff

## Abstract

Background compensation is the ability of a controlled variable to respond to an applied perturbation in an unchanged manner and independent of different but constant background signals which act in parallel to the perturbation. We found that background compensation occurs by ‘coherent feedback’ mechanisms where additional control variables feed directly back to the controlled variable. This paper extends a previous study on background compensation to include phase responses in frequency controlled coherent feedback oscillators. While the frequency resetting amplitude in coherent feedback oscillators is found to be dependent on the inflow/outflow perturbation of the controlled variable and thereby become phase dependent, the frequency resetting itself and the corresponding phase response curves are found to be background compensated. It is speculated that this type of background compensation may be an additional way how ambient noise can be ‘ignored’ by organisms.

## Introduction

Homeostatic mechanisms play an essential role in the defense of organisms against environmental and internal disturbances and thereby contribute to their stability. The term ‘homeostasis’ was introduced in 1929 by Walter Cannon [1, 2] similar to Claude Bernard’s concept [3] of the constancy of the internal milieu [4]. With the rise of cybernetics, Norbert Wiener related homeostasis to negative feedback mechanisms as ‘exemplified in mechanical automata’ [5]. In the following years Wiener’s negative feedback concept of homeostasis was applied to various physiological examples [6].

However, by the end of the 1980’s researchers became more critical that homeostasis would only relate to single negative feedback loops with a fixed setpoint. Focussing on properties such as variable setpoints, multiple feedbacks, the application to circadian rhythms, or on nonlinear dynamic behaviors alternative terms like ‘predictive homeostasis’ [7], ‘allostasis’ [8, 9], ‘rheostasis’ [10] or ‘homeodynamics’ [11, 12] were suggested instead and in addition [13] to ‘homeostasis’. As pointed out by Carpenter [14], although homeostatic regulation shows many different facets the term ‘homeostasis’ still can serve as an overarching concept.

In this paper I revisit theoretical work on oscillatory homeostats with robust frequency control. Frequency control has been observed in many biological systems, for example in circadian rhythms [15] or neuronal oscillations [16, 17]. While our understanding how frequency control is achieved in these systems is still relatively poor, I feel that insights into possible mechanisms of robust frequency and response control may be helpful to uncover principles behind such regulations.

### Aim of this work

In a previous study [18] we found that coherent feedback oscillators not only enable robust frequency homeostasis, but can also compensate for different but constant background perturbations. I extend here these earlier findings to include both inflow and outflow perturbations at the controlled variable, which show distinct phase dependencies in the oscillators’ frequency resetting. Despite these phase dependencies, the frequency resetting and the corresponding phase response curves (PRCs) turn out to be background compensated, while in ordinary single-loop feedback oscillators background compensation of the PRCs is not present. At the end of the paper I speculate whether background compensation may be an additional tool how organisms cope with ambient noise.

## Materials and methods

### Computational methods

Computations were performed with the Fortran subroutine LSODE [19] (https://computing.llnl.gov/projects/odepack). Plots were generated with gnuplot (www.gnuplot.info) and movies were made from a sequence of plots using QuickTime (https://support.apple.com/en-us/docs/software). Reaction schemes and plot annotations were prepared with Adobe Illustrator (www.adobe.com). Concentrations of compounds such as *A, E, I*_1_,… are described by their compound names without square brackets. Rate constants and other parameters are in arbitrary units (au) and represented by *k*_1_, *k*_2_, *k*_3_,… independent of their kinetic meaning, i.e. whether they are turnover numbers, Michaelis constants, or inhibition constants.

For documentation a set of selected computations are made available as Python and Matlab scripts (see supporting material S1 Programs).

### Phase response curves

We have used phase response curves to study the resetting dynamics of step- and pulse-perturbed oscillators. Phase responses have extensively been used in biology, especially in the study of circadian rhythms [20–23], but also in mechanistic analyses of purely chemical oscillators [24–26]. Fig 1 illustrates the method to determine a phase response curve. A perturbation (step or pulse) is applied at a certain phase and the resulting train of oscillations, outlined in blue, is compared with the corresponding undisturbed oscillator, which is outlined in gray (panel a). In a phase response curve (Fig 1b) the phase difference ΔΦ between corresponding peaks of perturbed and unperturbed oscillations is plotted against the phase of perturbation.

**Fig 1.**
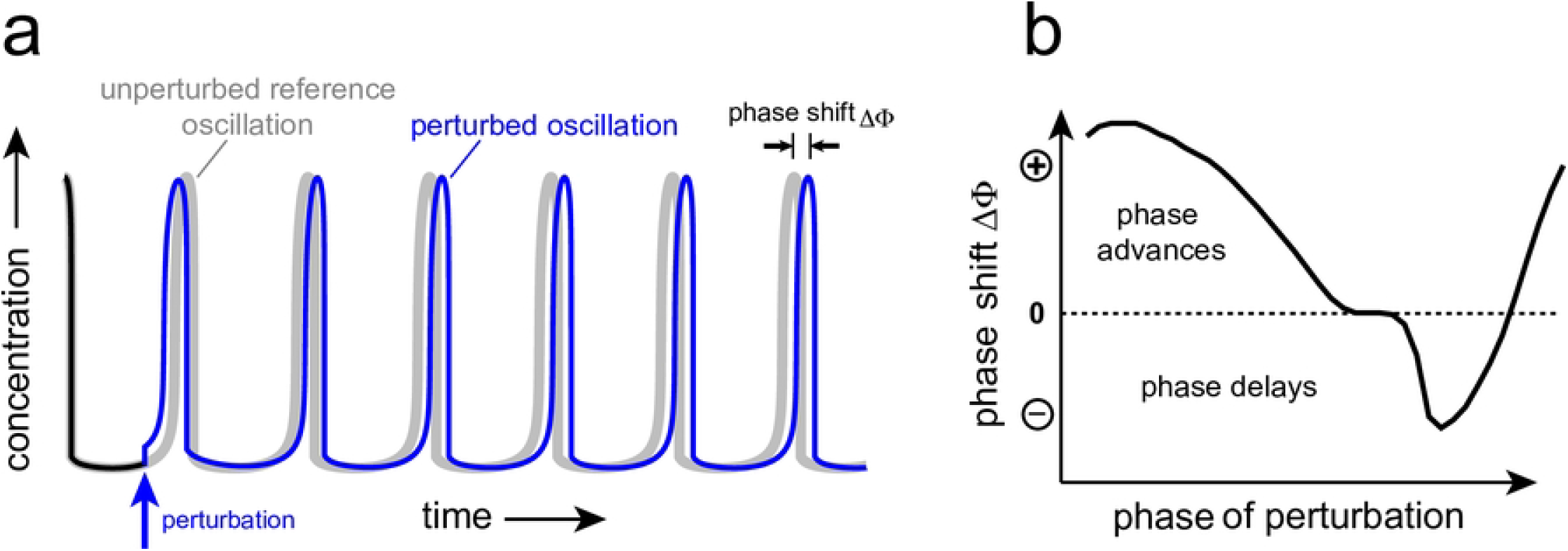
Determination of a phase response curve. Panel a: The application of a perturbation (indicated by the blue arrow) causes a phase shift ΔΦ between corresponding maxima of the perturbed and unperturbed oscillations. Panel b: A phase response curve is constructed by plotting phase shifts against the phases of perturbations, which are applied within one cycle of the unperturbed oscillation.

In biology positive phase shifts are generally related to *phase advances*, while negative phase shifts relate to *phase delays*, which leads to the definition of ΔΦ as

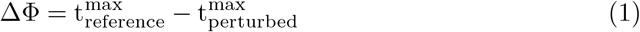

where 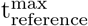 is the time of a maximum of the unperturbed oscillation, while 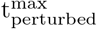 is the time of the corresponding maximum after the perturbation has been applied.

### Calculating averages

Averages of an oscillatory compound *X* are described as *<X>* and have been calculated by two methods. In one method *X* is integrated for a given time period *τ* and the average is determined as the ratio between the integral of *X* and *τ*, i.e.:

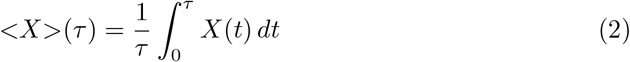

However, this method depends on the history of integrated *X*, which sometimes, especially after a rapid change in *X*, has the disadvantage that *<X>*(*τ*) may only slowly converge to its true value at *τ*.

In the other method I have used a self-chosen number (*N*_*sw*_) of overlapping time intervals or ‘sliding windows’ with time length Δ*t*, i.e. [*t*_*i*_, *t*_*i*_+Δ*t*]. The averages of *X* within each single window *i* is

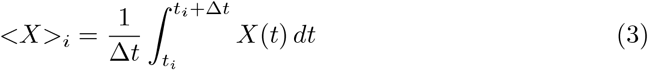

The *t*_*i*_’s are equal to LSODE’s step length and represent the successive time points during LSODE’s numerical integration. For this ‘sliding window method’ *<X>* is calculated as:

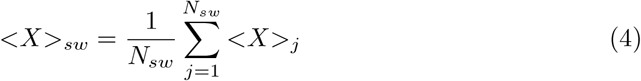

As a convention, *<X>*_*sw*_ is placed in the *X*-time plot at the time-middle of the last sliding window when *j* = *N*_*sw*_.

### Usage of zero-order kinetics

The reaction schemes I use include zero-order kinetics for two reasons: firstly, to introduce robust perfect adaptation in the controlled variables (see below), and secondly, to promote oscillations. Concerning the promotion of oscillations, Goodwin [27] presented in 1963 a two-variable negative feedback oscillator which since has been the basis for many physiological model oscillators [28]. An essential aspect in Goodwin’s 1963 oscillator is the presence of zero-order degradation of the two components. Thorsen et al. [29] showed that for conservative two-component negative feedback oscillators the above zero-order assumption leads to a general equation of the form

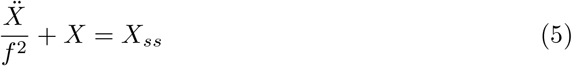

where *X* can be either of the two feedback variables, *X*_*ss*_ is the steady-state expression of *X*, where *f* approximately describes the frequency of the conservative oscillator (for details see Supporting Information in Ref [29]). In this respect, the conservative oscillator schemes which arise by the zero-order conditions can be viewed as a driving force for oscillations even when intermediates are present within the feedback loop.

Results along similar lines were also found by Kurosawa and Iwasa [30], who observed that introducing Michaelis-Menten kinetics in the degradation reactions of circadian clock models promote oscillations.

## Results and discussion

### Outline

First I will describe the properties and breakdown conditions of two single-loop feedback oscillators based on motif 2 and motif 8 (for overview of the feedback motifs, see [31]). Then, two additional controllers (*I*_1_ and *I*_2_) will be added to each of the single-loop feedbacks such that *I*_1_ and *I*_2_ will feed directly back to the controlled variable (*A*) and thereby define a coherent feedback loop structure [18], which is found to rescue the oscillators from their breakdown conditions. Then I show that the coherent feedbacks lead to background compensation in their frequency resetting for *both* inflow or outflow step perturbations. Finally I demonstrate that the resulting PRCs are background independent, followed by a brief summary and discussion.

### Motif 2 based controllers

#### Integral control of *A* oscillations

Fig 2 shows a single-loop feedback scheme which is related to Goodwin’s 1963 oscillator, except for the additional variable *e*.

**Fig 2.**
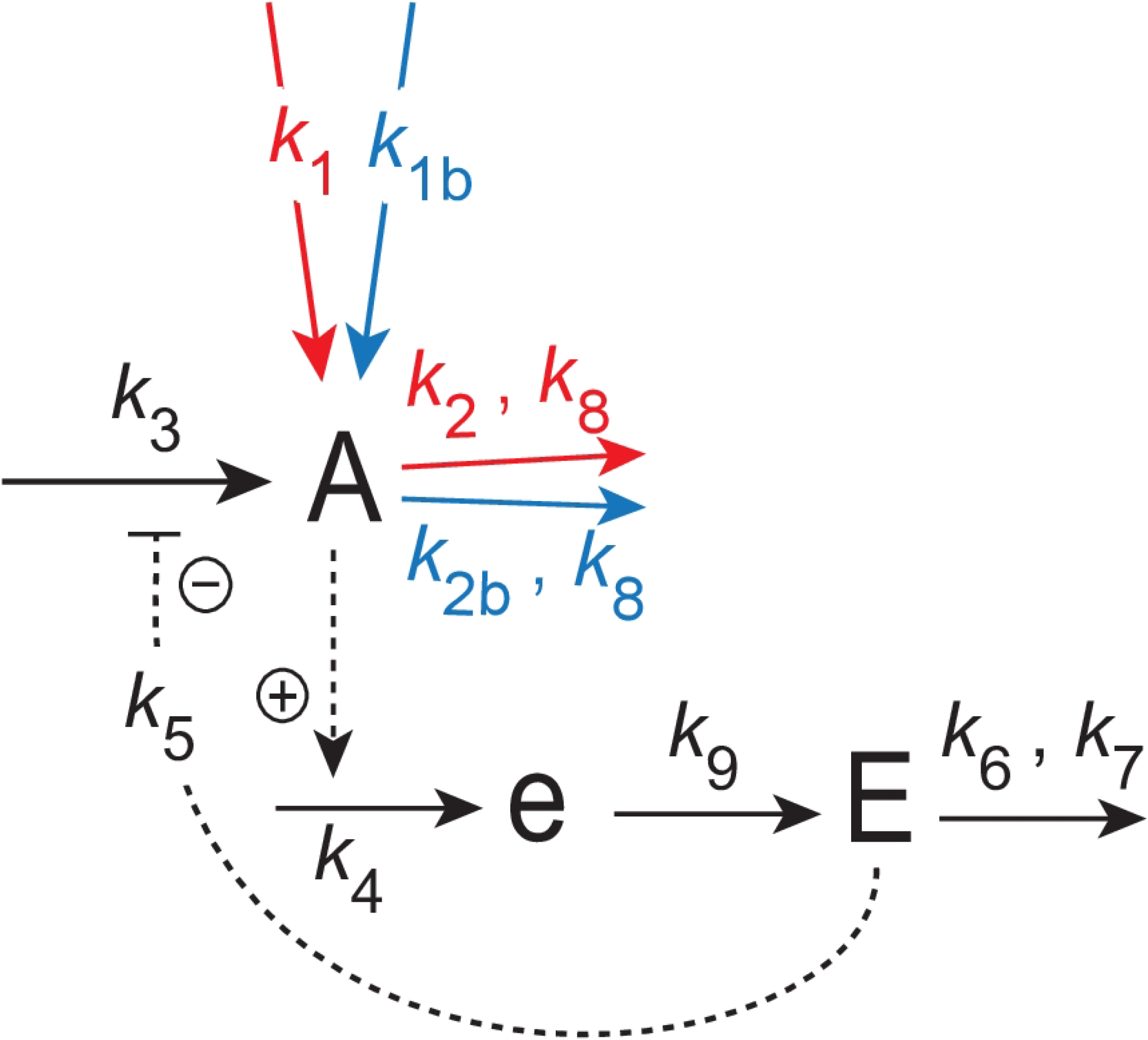
Feedback arrangement of a single-loop feedback similar to Goodwin’s 1963 oscillator, which is based on motif 2 [31]. Compound *A* is the controlled variable, while *E* is the controller. Compound *e* is an intermediate which assures limit cycle oscillations when the system is oscillatory. Red reaction arrows indicate applied step perturbations. The blue arrows indicate constant backgrounds.

The rate equations are:

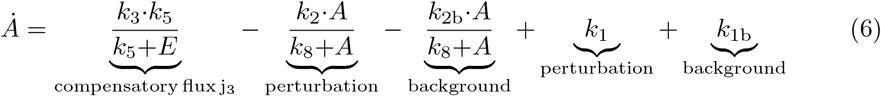

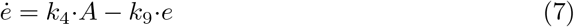

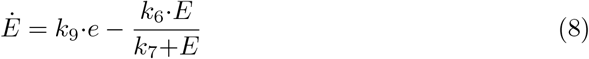

Dependent on the perturbation on *A* variable *E* controls *A* by acting either as a repressor or derepressor thereby adjusting the compensatory flux *j*_3_=*k*_3_*k*_5_*/*(*k*_5_+*E*). To obtain integral control in variable *A* (or *<A>* when the system is oscillatory) the difference between the setpoint of *A* or *<A>* and its actual value is calculated, which is integrated in time. The integrated error is then used to correct for an applied perturbation in *A* [32–35]. There are multiple kinetic conditions which can result in integral control [36–40]. The approach taken here uses zero-order kinetics [31, 41–44] with the interpretation [31, 43] that certain control-related enzymes work under saturated or near saturation condition. Another approach that can be used is based on a second-order removal of two variables (‘antithetic control’) [39, 45, 46]. Robust perfect adaptation can also be achieved by a first-order autocatalytic synthesis of the controller variable combined with its first-order degradation [47–49].

Since the scheme in Fig 2 is forced to be oscillatory by zero-order removals of *A* and *E* we assume that the average concentrations in *A, E*, and *e* are at steady state, i.e. 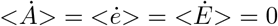. The setpoint of the controlled variable is *k*_6_*/k*_4_, which is obtained by combining Eqs 7 and 8 and eliminating the term *k*_9_ *e*. Taking then the average and solving for *<A>* we get:

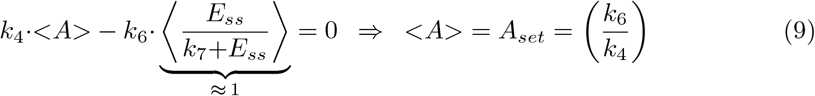

The feedback motif 2 in Fig 2 is what has been termed an *inflow controller* [31], which points to the fact that the compensatory flux *j*_3_ = *k*_3_*k*_5_*/*(*k*_5_+*E*) is an *inflow* to the controlled variable *A* (compensating outflows). The range of *k*_2_ for which the feedback can show homeostatic control in *<A>* is limited by the following two conditions:

i. by the maximum average compensatory flux *<j*_3_*>*=*k*_3_, which is reached when 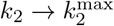 and *E* → 0 with

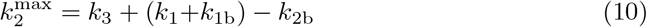

from Eq 6. Controller breakdown occurs when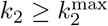.
ii. by the minimum compensatory flux *<j*_3_*>*=0, which is reached when the total inflow to *A* balances or becomes larger than the total outflow from *A*, i.e.

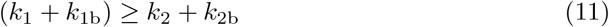

In this case *A* and *E* increase spontaneously (termed windup) while the system tries to satisfy the formal relationship *j*_3_=*k*_3_*k*_5_*/*(*k*_3_+*E*)=0. Fig 3 illustrates this type of controller breakdown by a successive increase of *k*_1_. As we will show below the limitation due to Eq 11 can be circumvented in a frequency-controlled version of the single-loop feedback when ‘outer control loop’ species *I*_1_ and *I*_2_ act as inflow and outflow controllers to both *A* and *E*.

**Fig 3.**
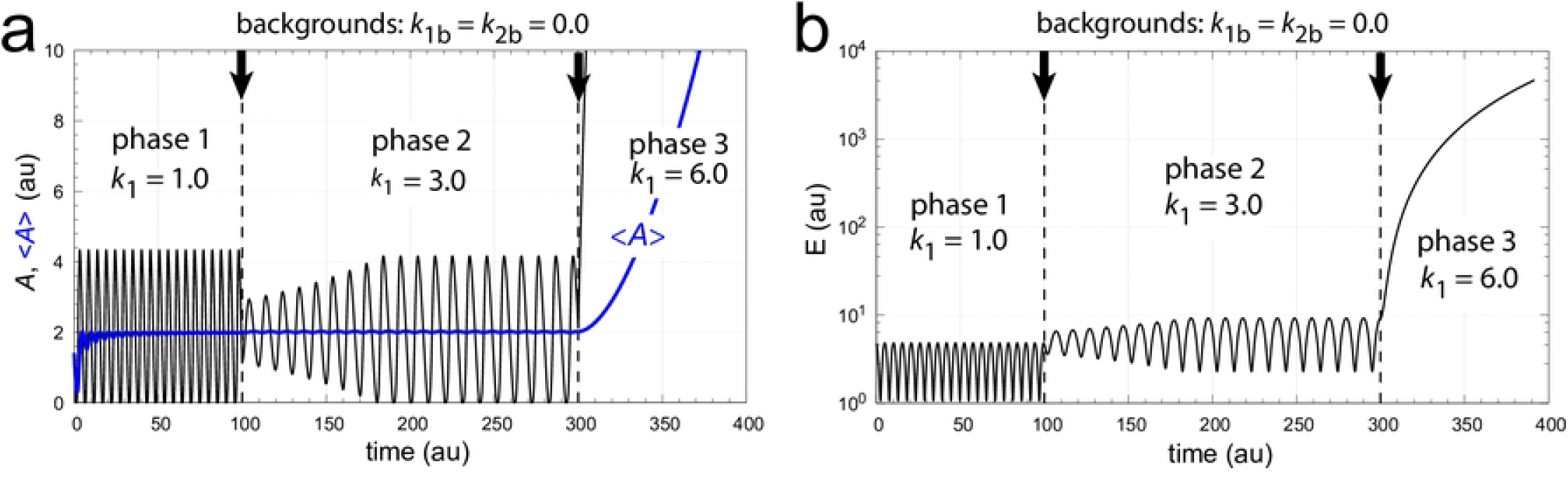
Illustration of controller breakdown of the scheme in Fig 2 when inflows to *A* exceed the outflows from *A*. The setpoint of *<A>* is 2.0. Panel a: *A* (in black) and *<A>* (in blue) are shown as a function of time when *k*_1_ increases successively from *k*_1_=1.0 (phase 1) to *k*_1_=3.0 (phase 2), and finally to *k*_1_=6.0 in phase 3. The final *k*_1_ satisfies Eq 11 which leads to controller breakdown and to a rapid increase in *A*. The average *<A>* is calculated after Eq 2. Vertical arrows show the times the *k*_1_ steps are increased. Also note the decrease in the oscillator’s frequency when *k*_1_ gets larger. Panel b: *E* as a function of time for the same *k*_1_ steps as in panel a. Frequency phase 1: 0.190; frequency phase 2: 0.097. Rate constants: *k*_1*b*_=0.0, *k*_2_=5.0, *k*_2*b*_=0.0, *k*_3_=100.0, *k*_4_=1.0, *k*_5_=0.1, *k*_6_=2.0, *k*_7_=*k*_8_=1×10^−6^, *k*_9_=20.0. Initial concentrations: *A*_0_=1.407, *E*_0_=4.754, *e*_0_=7.524×10^−2^.

There is also a third condition of controller breakdown when one of the signaling events (indicated by the dashed lines in Fig 2) reach saturation [44]. For the sake of simplicity we have not considered this possibility here and assumed that the activation of *e* by *A* is described by first-order kinetics with respect to the activator, i.e. *j*_4_=*k*_4_ *A* without an activation constant.

#### Introducing coherent feedback and frequency control

An interesting feedback arrangement occurs when additional controllers *I*_1_ and *I*_2_ in Fig 4 feed directly back to *A*. In this case robust homeostasis in both *A* and *E* can be achieved. The control of *E* by *I*_1_ and *I*_2_ leads in addition to frequency homeostasis [29]. In analogy to a similar feedback arrangement in quantum control theory [50, 51] we have termed the feedback scheme in Fig 4 as ‘coherent feedback’ [18].

**Fig 4.**
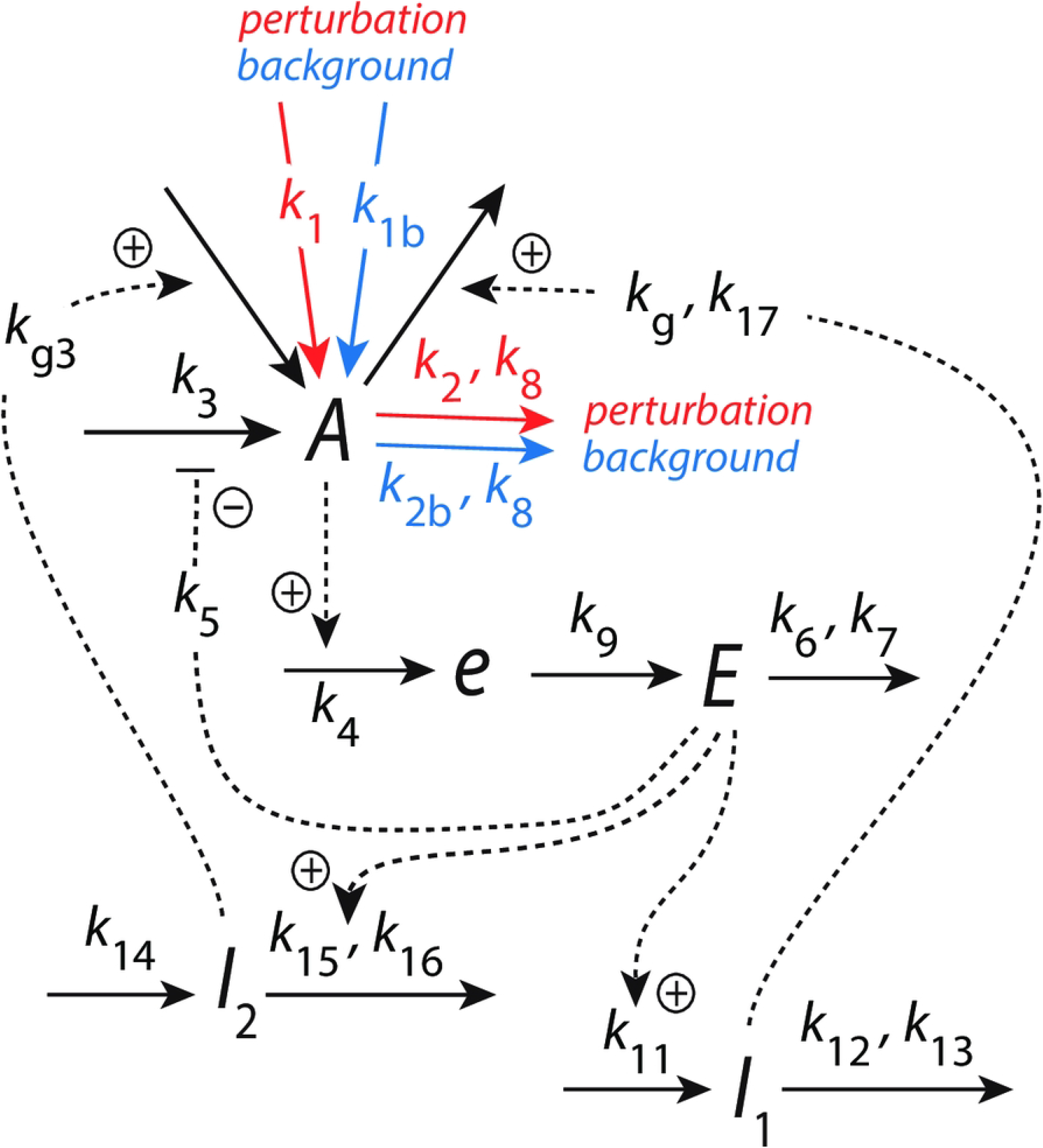
Coherent feedback keeps *<A>, <E>* and the frequency under homeostatic control by *I*_1_ and *I*_2_, which directly feed back to variable *A*.

The rate equations are:

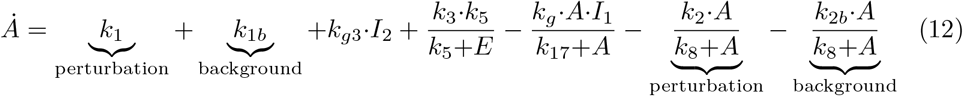

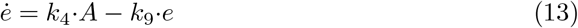

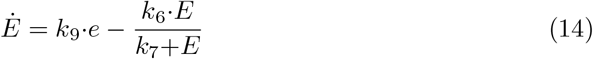

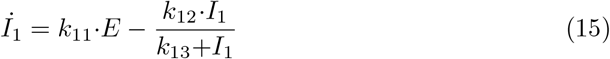

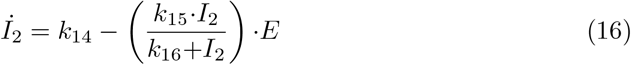

*k*_1_ or *k*_2_ denote step perturbations while *k*_1*b*_ and *k*_2*b*_ represent constant backgrounds. The setpoint of *<A>* is still given by Eq 9, while two setpoints for *<E>* are obtained from the rate equations of respectively *I*_1_ and *I*_2_, i.e. [18, 29]

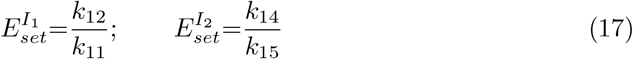

For the sake of simplicity we here keep both 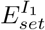 and 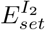 rather arbitrary equal to 5.0 and *A*_*set*_ to 2.0.

#### Background compensation in frequency resetting of M2 feedback loops

The interesting aspect of the coherent feedback scheme in Fig 4 is the fact that it can compensate for different but constant background perturbations. In our previous work [18] only *outflow* perturbations were considered, because the central feedback loop *A*-*e*-*E*-*A* in Fig 4 (being an *inflow* controller) compensates essentially for *outflow* perturbations in *A* [31]. However, since the *I*_1_-*I*_2_ ‘outer feedback layer’ in Fig 4 should also allow to compensate for inflows to *A* as implied by the work of Thorsen [52], I tested the coherent feedback scheme with respect to inflow perturbations to *A*. In Fig 5 we have the same rate constants and perturbative conditions as in Fig 3, but keep both *A* and *E* under homeostatic control by *I*_1_ and *I*_2_ (see Figs 5a and b). Fig 5c shows that the increasing *k*_1_ steps lead to an increase of *I*_1_ and a decrease of *I*_2_ which both contribute to the compensation of the inflow to *A*. Fig 5d shows that the steady state frequency is independent of *k*_1_.

**Fig 5.**
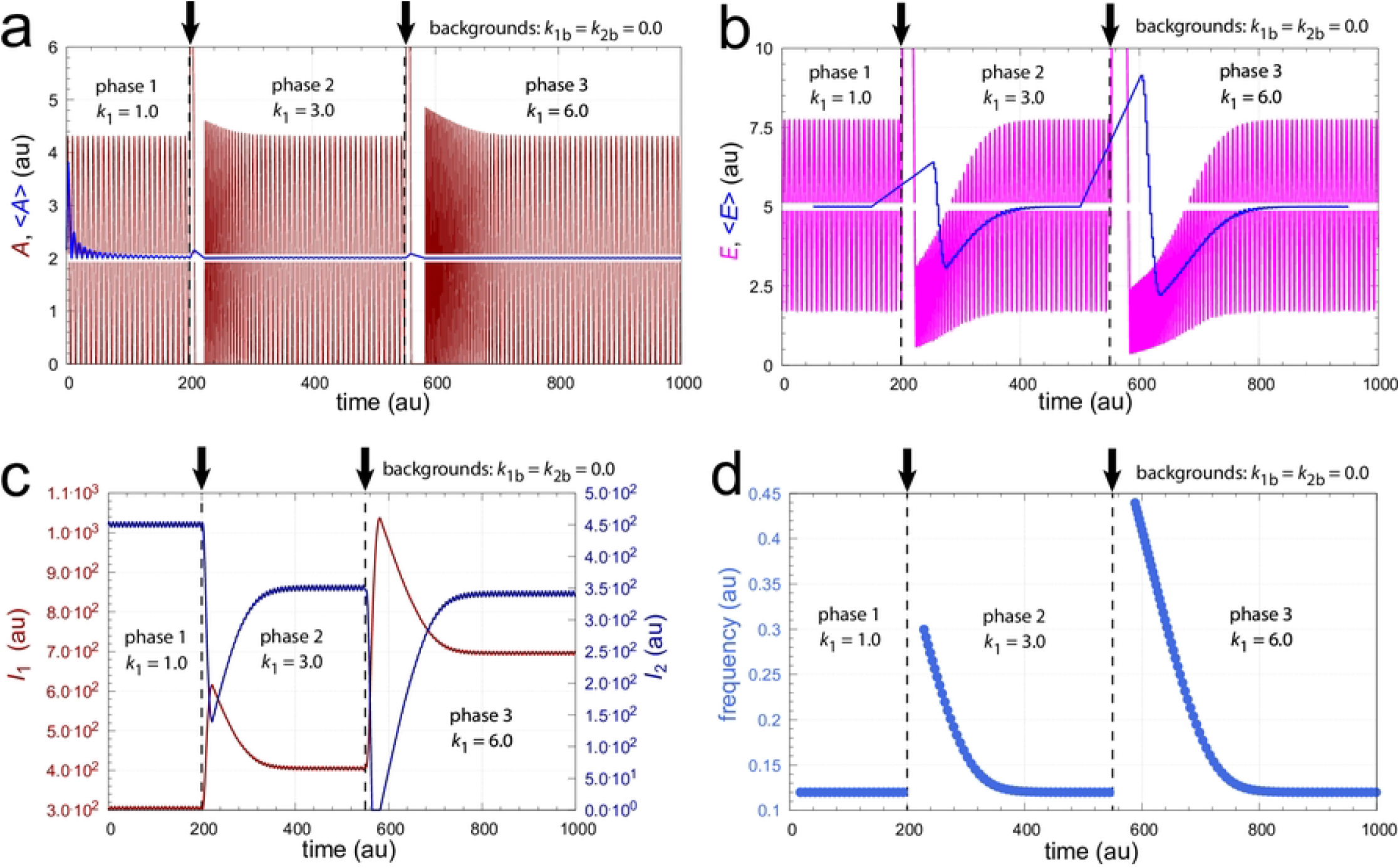
Compensation of inflow perturbations to *A* by controllers *I*_1_ and *I*_2_. Panel a: *A* and *<A>* are shown as a function of time. The setpoint of *A* is 2.0. Blue line shows *<A>* calculated by Eq 2. Panel b: *E* and *<E>* as a function of time. The setpoint of *E* is 5.0. Blue line shows *<E>* calculated by Eqs 3 and 4 with Δ*t* (sliding window size)=100.0 time units, *N*_*sw*_=50, and step length=0.05. Panel c: *I*_1_ and *I*_2_ as a function of time. Panel d: Frequency as a function of time. Vertical arrows indicate the time points when the *k*_1_ steps occur. Rate constants: *k*_1*b*_=0.0, *k*_2_=5.0, *k*_2*b*_=0.0, *k*_3_=100.0, *k*_4_=1.0, *k*_5_=0.1, *k*_6_=2.0, *k*_7_=*k*_8_=1×10^−6^, *k*_9_=20.0, *k*_11_=1.0, *k*_12_=5.0, *k*_13_=1×10^−6^, *k*_14_=5.0, *k*_15_=1.0, *k*_16_=*k*_17_=1×10^−6^, *k*_*g*_=*k*_*g*3_=0.01. Initial concentrations: *A*_0_=2.080, *E*_0_=1.731, *e*_0_=9.677×10^−2^, *I*_1,0_=304.87, *I*_2,0_=450.57.

Since the scheme in Fig 4 is stable against inflows and outflows to and from *A* I have tested its frequency resetting behavior for step perturbations by both *k*_1_ and *k*_2_. Fig 6 shows surprising differences in the frequency resetting: when *A* is perturbed by *k*_1_ steps the frequency resetting is highly dependent on the phase of the perturbation (Fig 6a). On the other hand, when *k*_2_ outflow steps are applied, the resetting of the frequency is practically independent of the phase where the perturbation is applied (Fig 6b).

**Fig 6.**
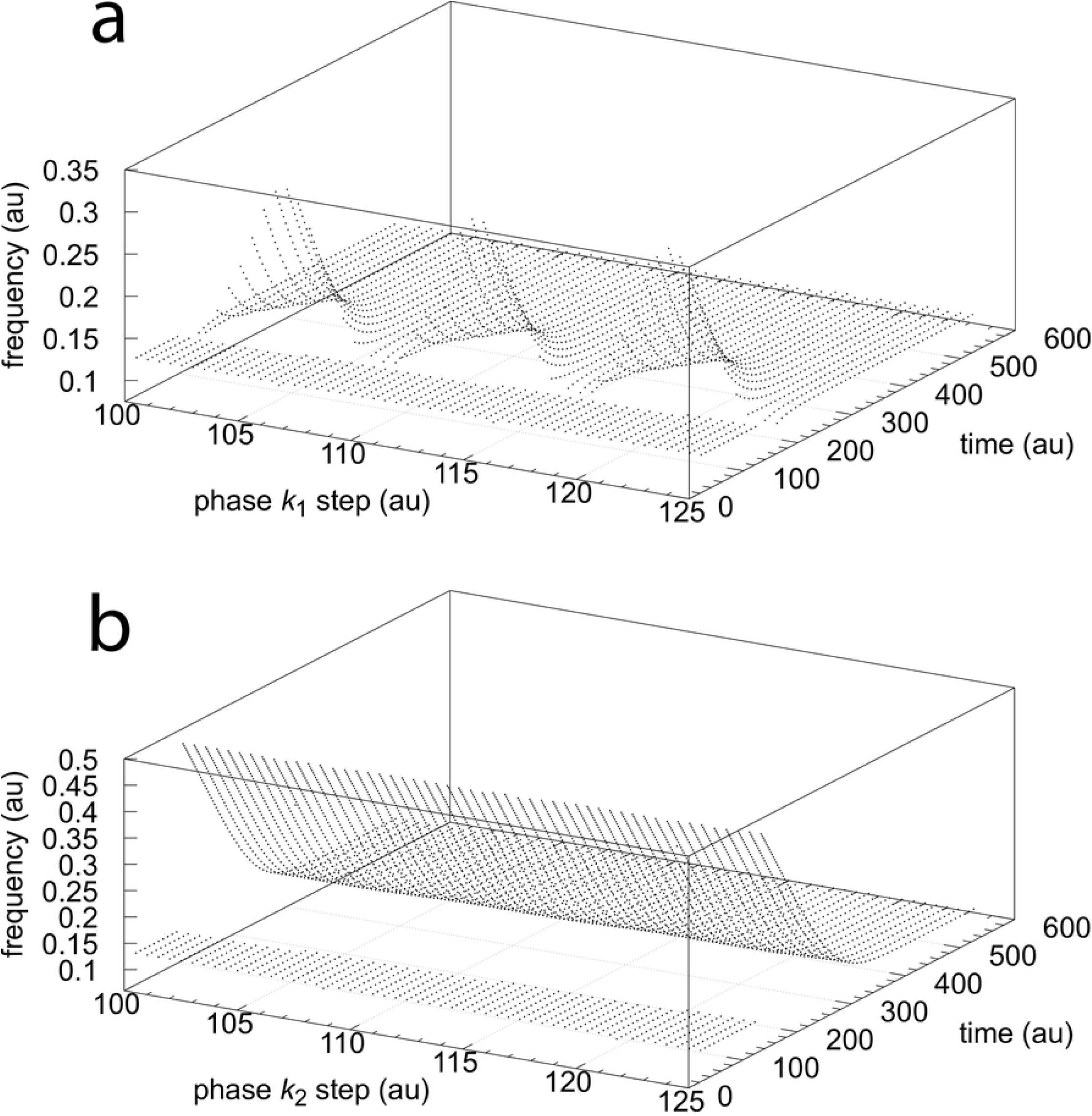
Phase dependencies of the frequency resetting in the oscillator Fig 4. Panel a: a *k*_1_ step perturbation of 1.0 → 3.0 is applied at different times/phases starting at t=100.0 and ending at t=125.0 with intervals of 0.5. At each phase the resetting frequency is plotted against time. Rate constants: *k*_1*b*_=20.0, *k*_2_=1.0, *k*_2*b*_=32.0, *k*_3_=100.0, *k*_4_=1.0, *k*_5_=0.1, *k*_6_=2.0, *k*_7_=*k*_8_=1×10^−6^, *k*_9_=20.0, *k*_11_=1.0, *k*_12_=5.0, *k*_13_=1×10^−6^, *k*_14_=5.0, *k*_15_=1.0, *k*_16_=*k*_17_=1×10^−6^, *k*_*g*_=*k*_*g*3_=0.01. Initial concentrations: *A*_0_=1.262, *E*_0_=7.550, *e*_0_=6.625×10^−2^, *I*_1,0_=2766.7, *I*_2,0_=3709.6. Panel b: As panel a, but a *k*_2_ step perturbation of 1→10 is applied with *k*_1_=1.0. Other rate constants, backgrounds *k*_1*b*_ and *k*_2*b*_, and initial concentrations as in panel a.

A closer look reveals that the difference in the two resetting behaviors is caused by the topology of the phase space kinetics. When *k*_1_ steps are applied the trajectory makes large excursions in phase space and returns to its oscillatory state after a transient. In case of *k*_2_ steps the system is rapidly pushed into an oscillatory state via a very short transit at low *A* concentrations along the *I*_1_-*E* or *I*_2_-*E* manifolds. Figs 7 and 8 show the time profiles of *A, E, I*_1_, *I*_2_, and the frequency when *k*_1_ and *k*_2_ steps are respectively applied at time *t* = 100. In both Fig 7 and Fig 8 the setpoints of *A*_set_=2.0 and *E*_set_=5.0 are defended due to the compensatory actions by *I*_1_ and *I*_2_, which results in the frequency homeostasis of the oscillator.

**Fig 7.**
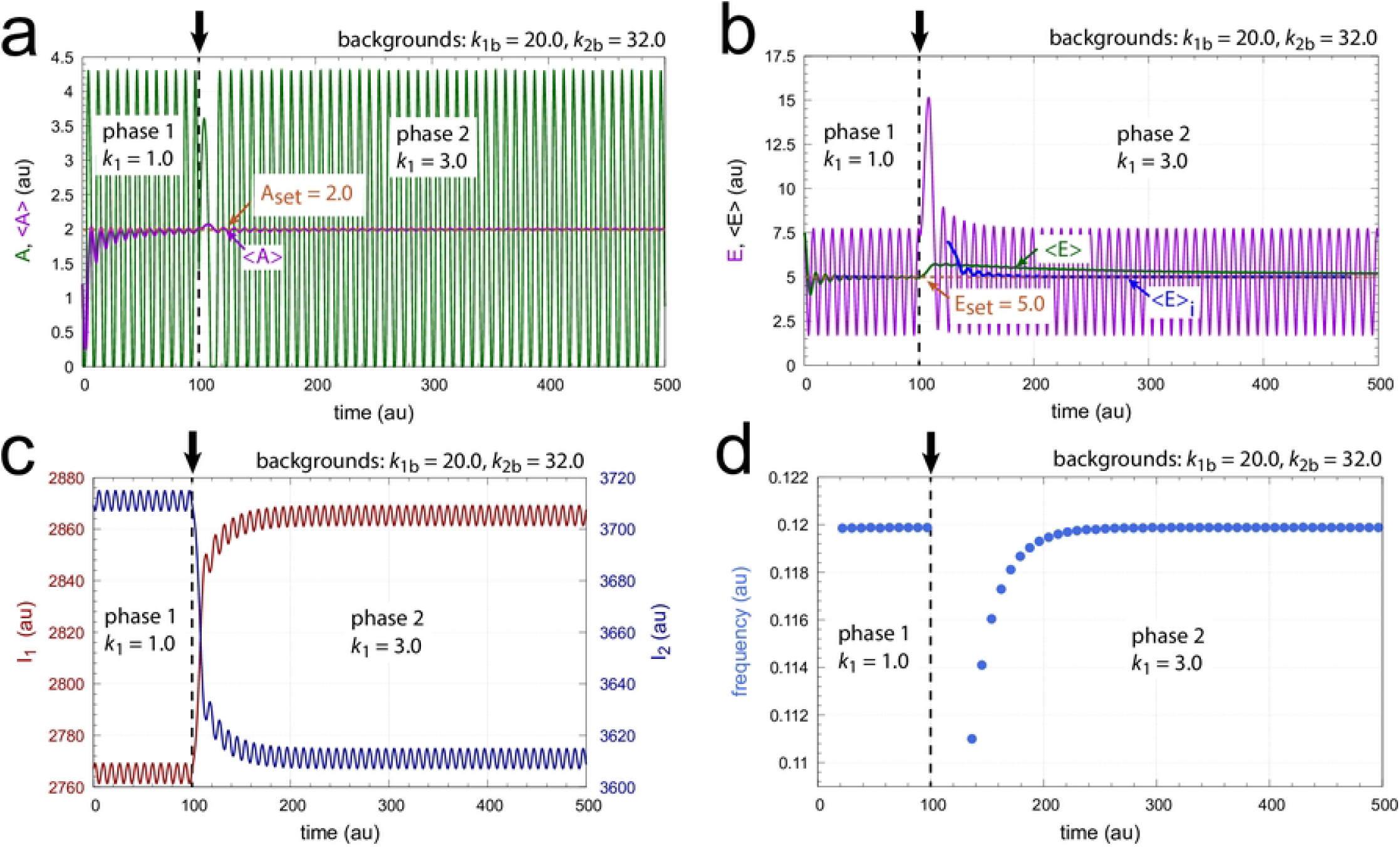
Time profiles of *A, E, I*_1_, *I*_2_, and the frequency when a *k*_1_ : 1.0 → 3.0 step is applied at time *t* = 100.0 (indicated by the vertical arrows). Panel a: Concentration of *A* as a function of time. Averages *<A>* are calculated by Eq 2. Panel b: Concentration of *E* as a function of time. Averages *<E>* (green line) are calculated by Eq 2 while *<E>*_i_ values (blue line) are calculated by Eq 3 using 500 time intervals of 50 units in phase 1 and 3500 intervals of the same extension in phase 2. Panel c: Concentrations of *I*_1_ and *I*_2_ as a function of time. Panel d: Frequency as a function of time. Rate constants and initial concentrations as in Fig 6.

**Fig 8.**
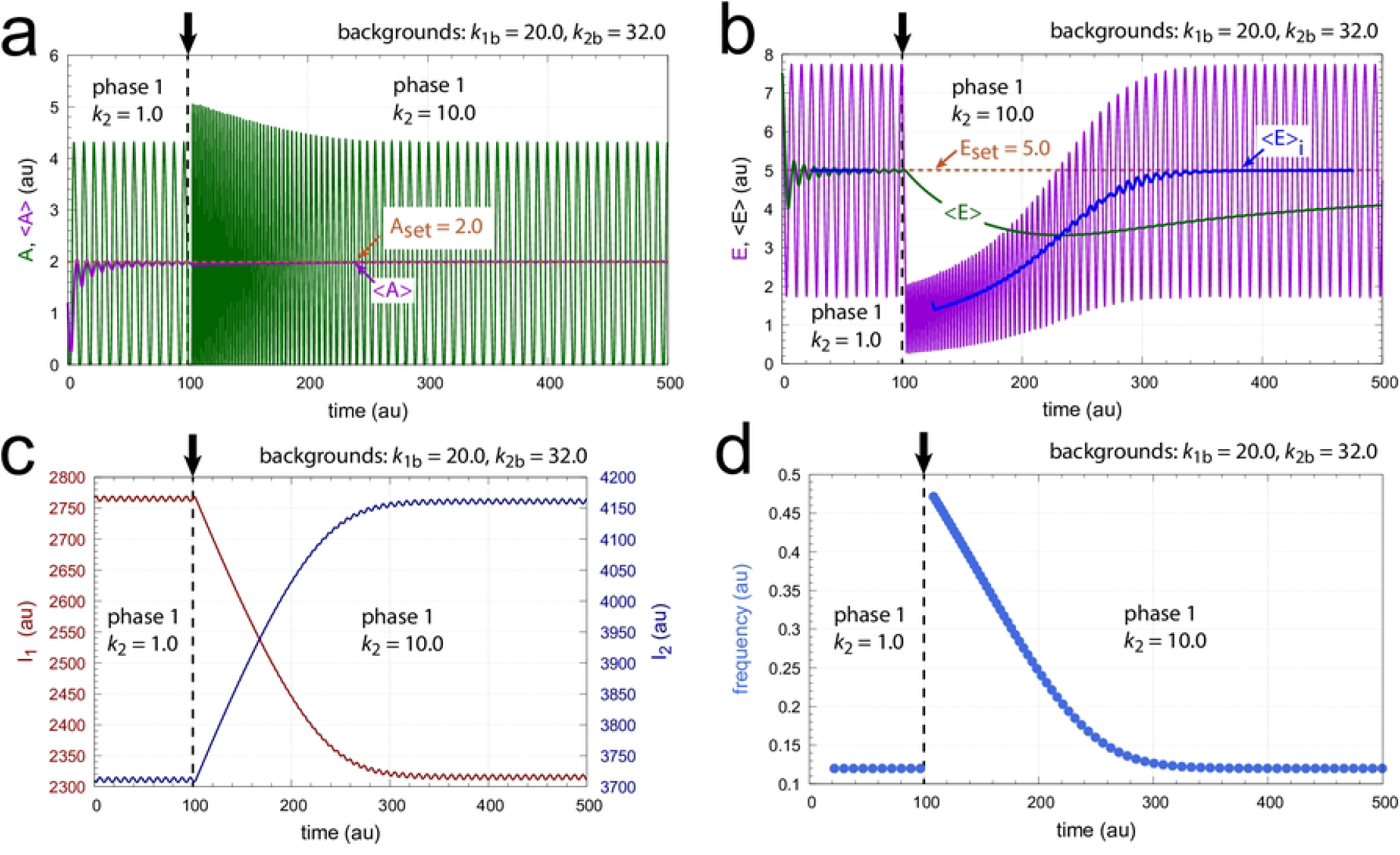
Time profiles of *A, E, I*_1_, *I*_2_, and the frequency when a *k*_2_ : 1.0 → 10.0 step is applied at time *t* = 100.0 (indicated by the vertical arrows) with *k*_1_=1.0. Other rate constants and initial concentrations as in Fig 6.

Fig 9 shows the trajectories of Figs 7 and 8 in *A*-*E*-*I*_1_ and *A*-*E*-*I*_2_ phase space.

**Fig 9.**
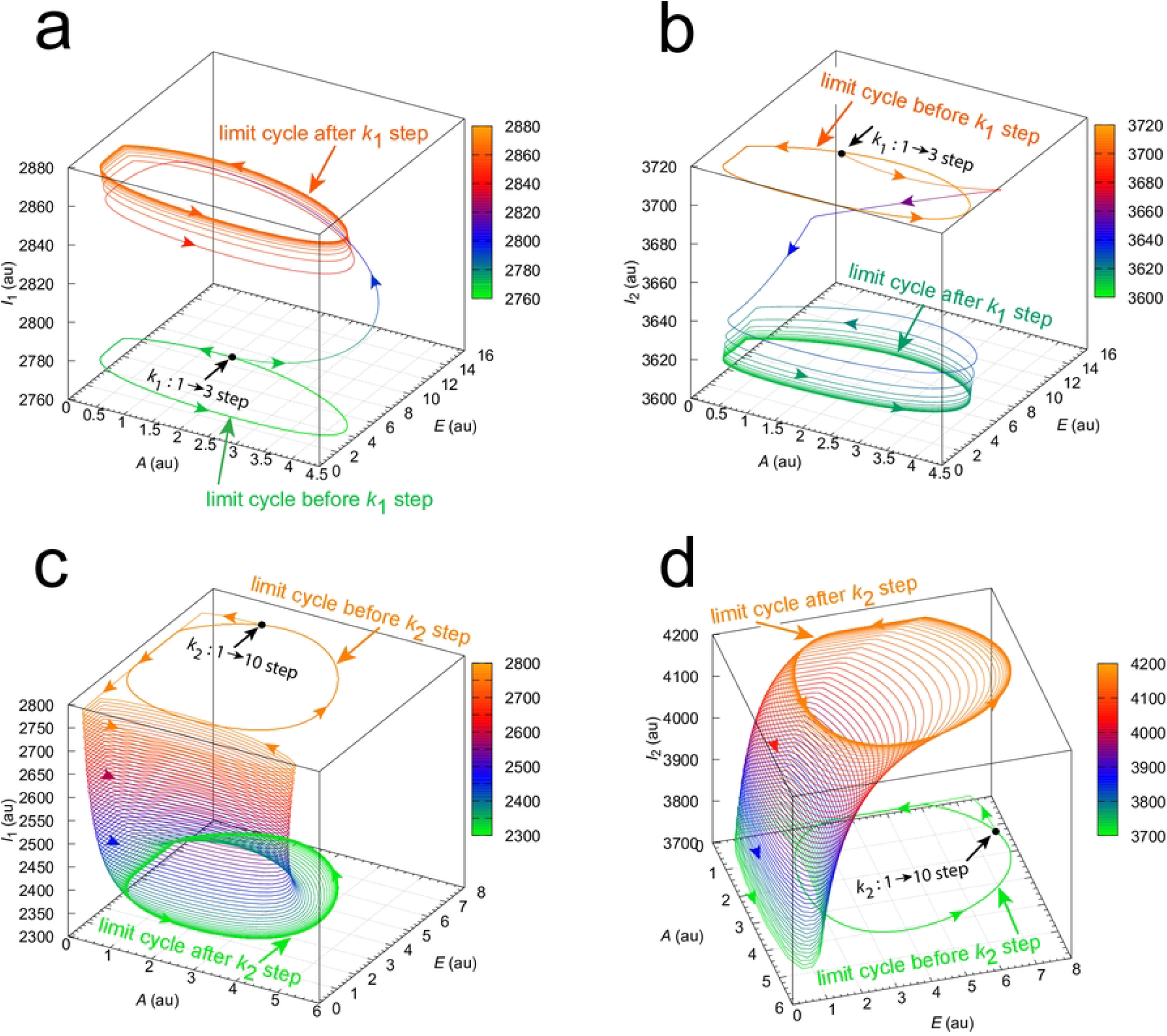
*A*-*E*-*I*_1_ and *A*-*E*-*I*_2_ phase space trajectories of the perturbed oscillators in Figs 7 and 8. Panels a and b show the trajectories for the *k*_1_ : 1.0 → 3.0 step at *t* = 100 in Fig 7. Panels c and d show the trajectories for the *k*_2_ : 1.0 → 10.0 step at *t* = 100 in Fig 8. Colors indicate the levels of *I*_1_ or *I*_2_.

The limit cycles projected on to the *A*-*E* phase space are preserved after the *k*_1_ or *k*_2_ steps. S1 Movies shows the trajectory excursions for the different resettings and the preservation of the limit cycles when projected on to the *A*-*E* phase space.

While Fig 9 provides insights into why the resetting of *k*_1_ steps is markedly different from those of *k*_2_ steps, in the next calculations I tested whether background compensation is operative for both *k*_1_ and *k*_2_ step perturbations. This is shown in Figs 10 and 11. As an example, I show two different applied backgrounds. In both figures the larger orange dots relate to *k*_1*b*_=20.0 and *k*_1*b*_=32.0, while the smaller blue dots refer to *k*_1*b*_=20.0 and *k*_1*b*_=40.0. For each time point a blue dot lies precisely at the position of a corresponding orange dot indicating that background compensation is operative. Supporting materials S2 Movie and S3 Movie show Figs 10a and 11a with varying viewing angles.

**Fig 10.**
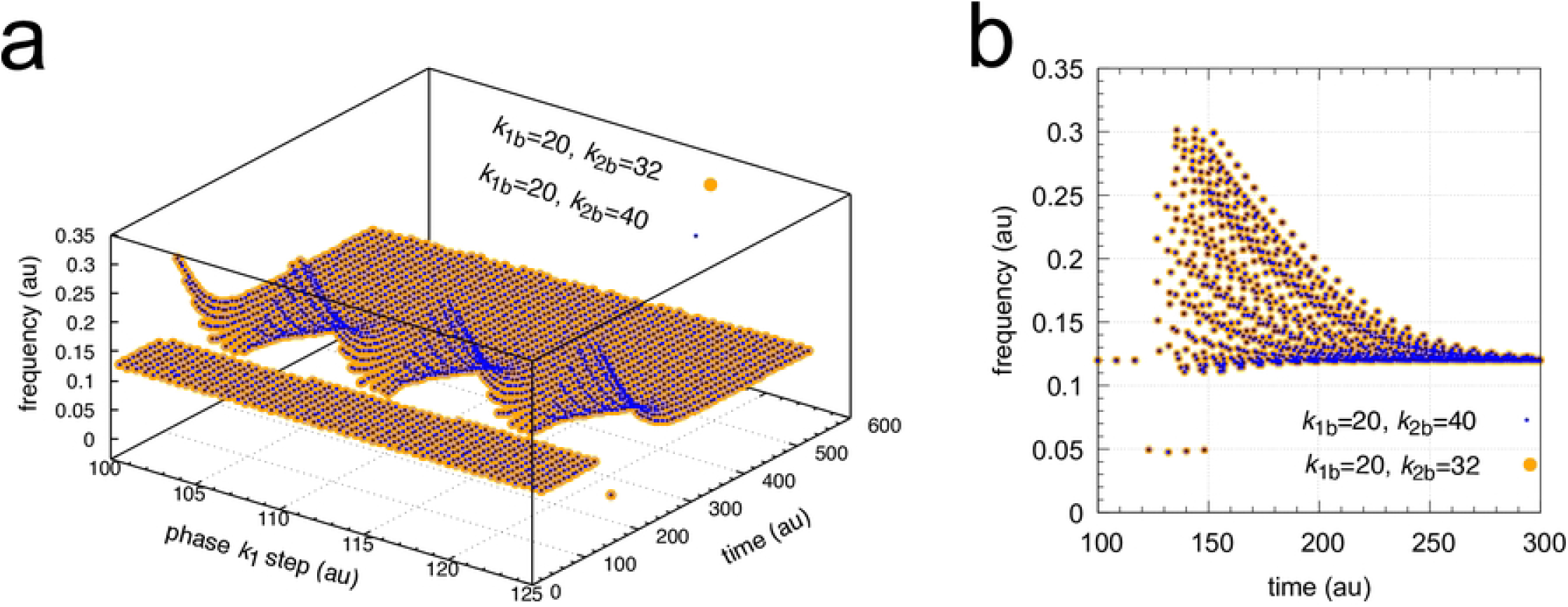
Background compensation in frequency resetting for *k*_1_ steps. *k*_1_ : 1.0 → 3.0 steps at two backgrounds are applied. Orange dots: *k*_1*b*_=20.0 and *k*_1*b*_=32.0; blue dots: *k*_1*b*_=20.0 and *k*_1*b*_=40.0. Panel a shows frequency as a function of phase of applied *k*_1_ steps and time, while panel b shows part of the projection on to the frequency-time axes. Rate constants as in Fig 6. Initial concentrations at *k*_1_ phase = 0: *A*_0_=4.304, *E*_0_=3.952, *e*_0_=2.150×10^−1^, *I*_1,0_=2761.6, *I*_2,0_=3714.7. See also S2 Movie.

**Fig 11.**
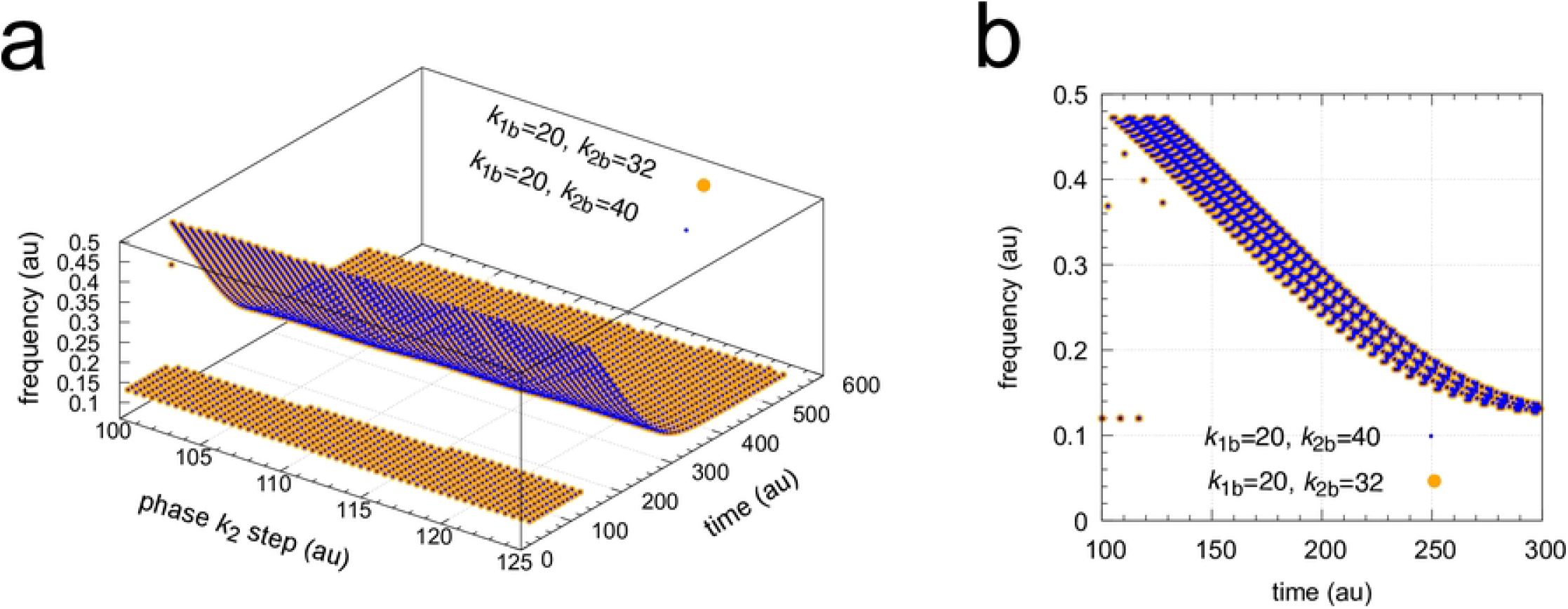
Background compensation in frequency resetting for *k*_2_ steps. *k*_2_ : 1.0 → 10.0 steps at two backgrounds are applied. Orange dots: *k*_1*b*_=20.0 and *k*_1*b*_=32.0; blue dots: *k*_1*b*_=20.0 and *k*_1*b*_=40.0. Panel a shows frequency as a function of phase of applied *k*_2_ steps and time, while panel b shows part of the projection on to the frequency-time axes. Rate constants as in Fig 6. Initial concentrations at *k*_2_ phase = 0 as in Fig 10. See also S3 Movie.

#### Background influences on PRCs in M2 feedback loops

Phase response curves (PRCs) are an often used tool to analyze biological or chemical oscillators. Especially have PRCs been used in relationship with circadian rhythms [20–23]. Due to the different phase resettings when *k*_1_ steps are applied to the oscillator in Fig 4 (see Figs 6a and 10a) I became interested to what extend phase shifts may be influenced by the applied backgrounds *k*_1*b*_ and *k*_2*b*_. Since increased backgrounds can lead to a diminished response amplitude in analogy to Weber’s law [53, 54], one may expect that Weber’s law may also apply to phase shifts. This expectation is, however, only partially fulfilled.

#### Single-loop M2 feedback

To investigate the influence of backgrounds on PRCs I first consider the single feedback loop in Fig 2 before turning to the coherent feedback scheme of Fig 4. A problem with the oscillator in Fig 2 is that the period will depend on the inflow/outflow rates to and from *A* (Fig 3) and thereby backgrounds have an influence on the oscillator’s frequency. In addition, phase shifts very often reach their final values first after a couple of cycles, which is illustrated in Fig 12. Fig 12a shows the application of a *k*_2_ pulse at phase *t*=1.0 leading to phase advances (positive phase shifts), while in panel b the pulse is applied at *t*=15.0 which leads to phase delays.

**Fig 12.**
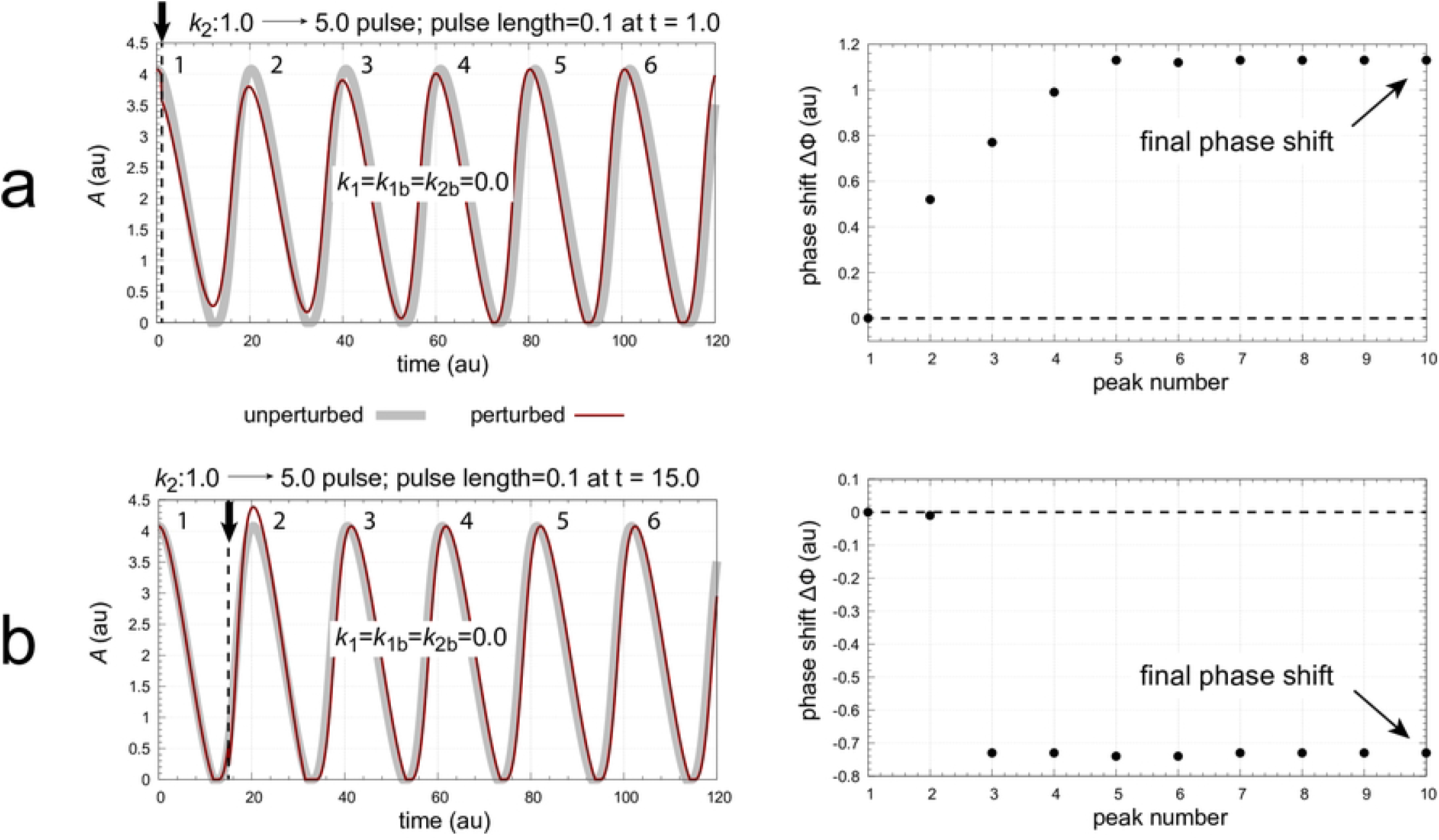
Positive and negative phase shifts are observed when *k*_2_ : 1.0 → 5.0 pulses of 0.1 time units are applied to the oscillator of Fig 2. Figures a and b, left panels: oscillatory concentrations of *A* for the unperturbed and perturbed system (outlined in gray and red) when the pulse is applied at respectively 1.0 and 15.0 time units. Peak numbers are indicated near peaks. Figures a and b, right panels: phase shifts as a function of the peak numbers from left panels. The unperturbed oscillator has a period length of 20.342 time units. Rate constants: *k*_1_=*k*_1*b*_=*k*_2*b*_=0.0, *k*_3_=100.0, *k*_4_=1.0, *k*_5_=0.1, *k*_6_=2.0, *k*_7_=*k*_8_=1×10^−6^, *k*_9_=20.0. Initial concentrations: *A*_0_=4.0763, *E*_0_=9.9288, and *e*_0_=0.2038.

The PRCs are constructed by plotting the final phase shifts (see Fig 12) against the phase of perturbation. To directly compare the PRCs at different backgrounds the ‘phase of perturbation’ is normalized with respect to the oscillator’s period length.

Phase=0 is defined to occur at an *A* maximum while the next maximum in *A* defines the normalized phase to be 1. Fig 13 shows the normalized phase response curves for different *k*_2*b*_ backgrounds when the same *k*_2_ pulse as in Fig 12 is applied. One sees a clear background dependency of the phase response curves: both the phase shift amplitude and the length of the dead zone (characterized by ΔΦ=0) increase with increasing *k*_2*b*_.

**Fig 13.**
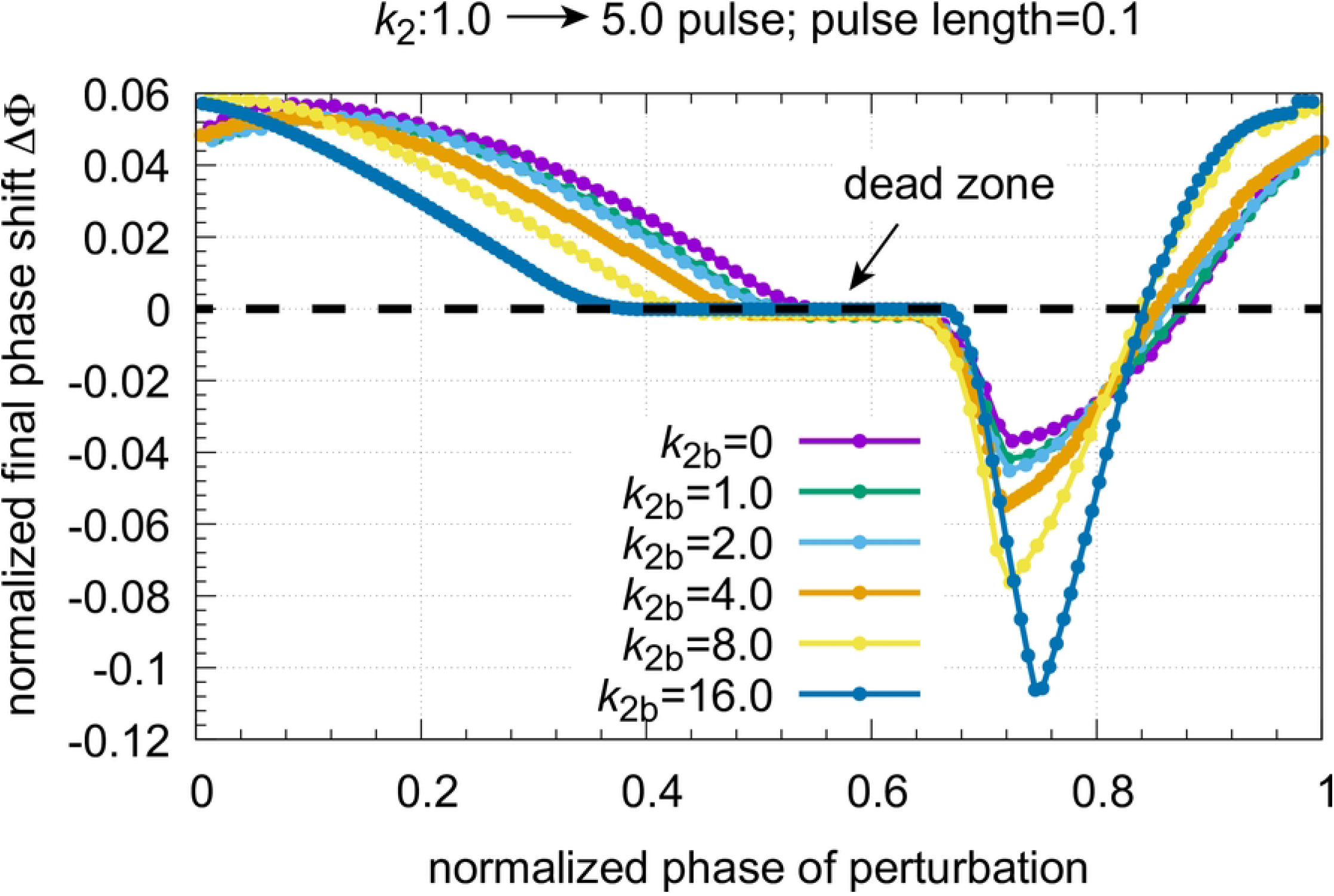
Normalized phase response curves at different *k*_2*b*_ backgrounds when a *k*_2_ : 1.0 → 5.0 pulse of 0.1 time length is applied to the oscillator of Fig 2. Initial concentrations, *k*_2*b*_=0.0: see legend Fig 12; initial concentrations *k*_2*b*_=1.0: *A*_0_=4.1570, *E*_0_=4.9277, *e*_0_=0.2078; period length: 10.265; initial concentrations *k*_2*b*_=2.0: *A*_0_=4.2416, *E*_0_=3.2540, and *e*_0_=0.2119, period length: 6.919; initial concentrations *k*_2*b*_=4.0: *A*_0_=4.4218, *E*_0_=1.9125, and *e*_0_=0.2381, period length: 4.263; initial concentrations *k*_2*b*_=8.0: *A*_0_=4.8168, *E*_0_=1.0398, and *e*_0_=0.2203, period length: 2.534; initial concentrations *k*_2*b*_=16.0: *A*_0_=5.6957, *E*_0_=0.4810, and *e*_0_=0.2684, period length: 1.583. Other rate constants as in Fig 12.

Concerning the dead zone, Uriu and Tei [55] recently found that saturation kinetics in the repressor synthesis appears responsible for the occurrence of a dead zone in circadian rhythms. However, in our case the repressor kinetics nor the compensatory flux *j*_3_ (see Eq 6) are saturated, but depend on the perturbations and backgrounds.

What is saturated in our model are the removals of *A* and *E*. This indicates that homeostatic constraints by zero-order kinetics may in addition lead to the appearance of a dead zone in the PRCs of circadian rhythms, an aspect which needs further investigations.

#### Coherent M2 feedback

Next I applied *k*_1_ : 1.0 → 128.0 and *k*_2_ : 1.0 → 128.0 step perturbations on the oscillator with coherent feedback (Fig 4) using the rather arbitrary chosen three backgrounds: (i) *k*_1*b*_=1.0 and *k*_1*b*_=500.0, (ii) *k*_1*b*_=1.0 and *k*_1*b*_=10.0, and (iii) *k*_1*b*_=100.0 and *k*_1*b*_=10.0. To illustrate the procedure, Fig 14 shows the resetting and the determined phase shifts when the phase of perturbation is 3.0 (panel a) and 4.4 (panel c) with backgrounds *k*_1*b*_=1.0 and *k*_1*b*_=500.0. Grayed oscillations in panels a and c represent the unperturbed rhythms, while the red oscillations show the effect of the perturbations. Panels b and d show the changes in phase shifts as a function of peak number and the final settling of the phase shift.

**Fig 14.**
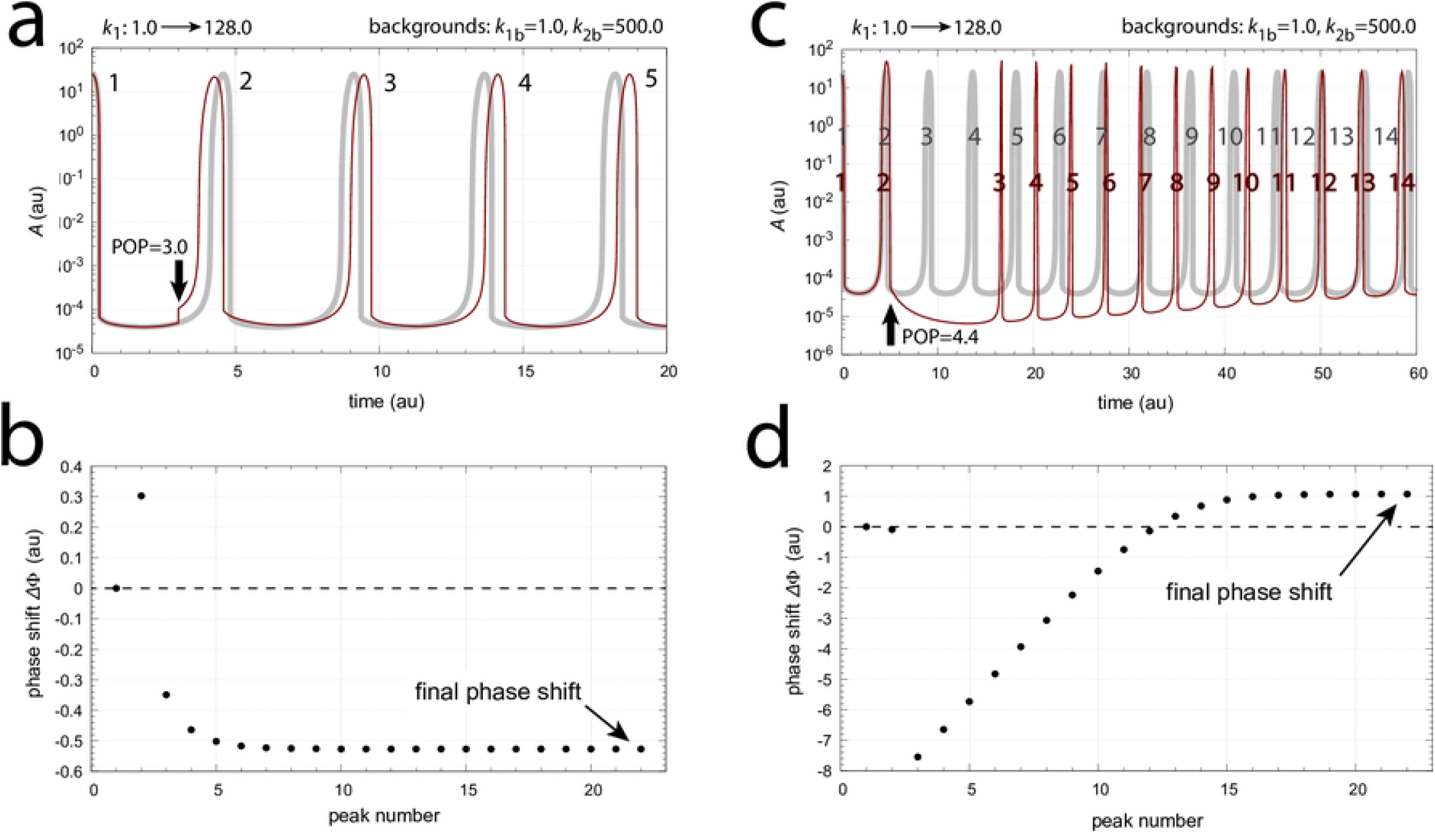
Two examples of the temporal behavior of phase shifts. Panel a: The applied phase of perturbation (POP) of a *k*_1_ : 1.0 → 128.0 step is 3.0 and indicated by the vertical arrow. The gray trace shows the unperturbed oscillation while the red trace shows the effect of the *k*_1_ step for the first five peaks. Numbers indicate the perturbed and unperturbed peaks. Panel b: Phase shift ΔΦ (Eq 1) as a function of peak number *i*. In total 22 peaks were recorded. Panel c: As panel a, but POP=4.4 as indicated by the vertical arrow. The picture shows the first fourteen peaks. Panel d: Phase shift ΔΦ (Eq 1) as a function of peak number *i*. Although phase shifts are in the beginning negative and reflect delays, the final phase shift is a phase advance. Rate constants: *k*_1*b*_=1.0, *k*_2_=1.0, *k*_2*b*_=500.0, *k*_3_=1×10^4^, *k*_4_=1.0, *k*_5_=0.1, *k*_6_=2.0, *k*_7_=*k*_8_=1×10^−6^, *k*_9_=2.0, *k*_11_=10.0, *k*_12_=50.0, *k*_13_=1×10^−6^, *k*_14_=50.0, *k*_15_=10.0, *k*_16_=*k*_17_=1×10^−6^, *k*_*g*_=*k*_*g*3_=1.0. Initial concentrations: *A*_0_=24.921, *E*_0_=2.433, *e*_0_=3.984, *I*_1,0_=1.423×10^4^, *I*_2,0_=1.153×10^4^.

Fig 14 shows the calculated phase response curves of *k*_1_ and *k*_2_ 1.0 → 128.0 steps for the three backgrounds (i)-(iii) indicated above. The PRCs are completely congruent, although slight differences in the final phase shifts are observed when phase shifts are outside of the constant phase shift zone. Surprisingly, this constant phase shift zone resembles that of a dead zone, but the final phase shift values are either negative (Fig 15a) or positive (Fig 15b). Other rate constants as in Fig 14.

**Fig 15.**
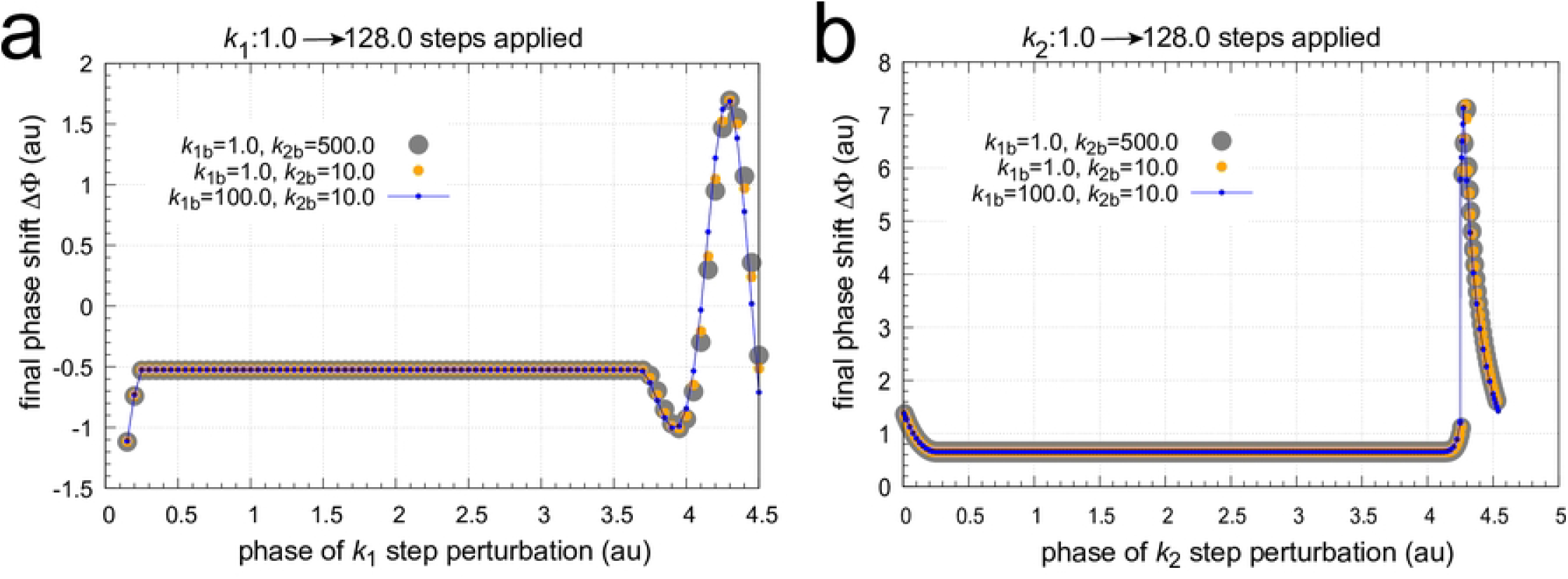
Phase response curves of the frequency compensated coherent feedback scheme in Fig 4 when *k*_1_ and *k*_2_ step perturbations are applied at three different backgrounds. Panel a: The phase response curve for *k*_1_ : 1.0 → 128.0 steps. Panel b: The phase response curve for *k*_2_ : 1.0 → 128.0 steps. Initial concentrations for the reference oscillations (*k*_1_=*k*_2_=1.0): (i) *k*_1*b*_=1.0, *k*_2*b*_=500.0: see caption Fig 14; (ii) *k*_1*b*_=1.0, *k*_2*b*_=10.0: *A*=24.839, *E*=2.447, *e*=4.000, *I*_1_=3.414 × 10^4^, *I*_2_=3.375 × 10^4^; (iii) *k*_1*b*_=100.0, *k*_2*b*_=10.0: *A*=24.839, *E*=2.442, *e*=3.987, *I*_1_=3.419 × 10^4^, *I*_2_=3.370 × 10^4^.

### Motif 8 based controllers

In the next set of calculations I show results for a Motif 8 (M8) feedback scheme [31], which shows flux control instead of concentration control.

#### M8 single-loop: Integral control of *A*-regulated flux and its breakdown by dominant outflow of *A*

Fig 16 shows the scheme of the considered M8 single negative feedback loop with *e* as a precursor of controller *E*. The role of the addition of *e* to the feedback is to turn an otherwise conservative system into a limit cycle oscillator.

**Fig 16.**
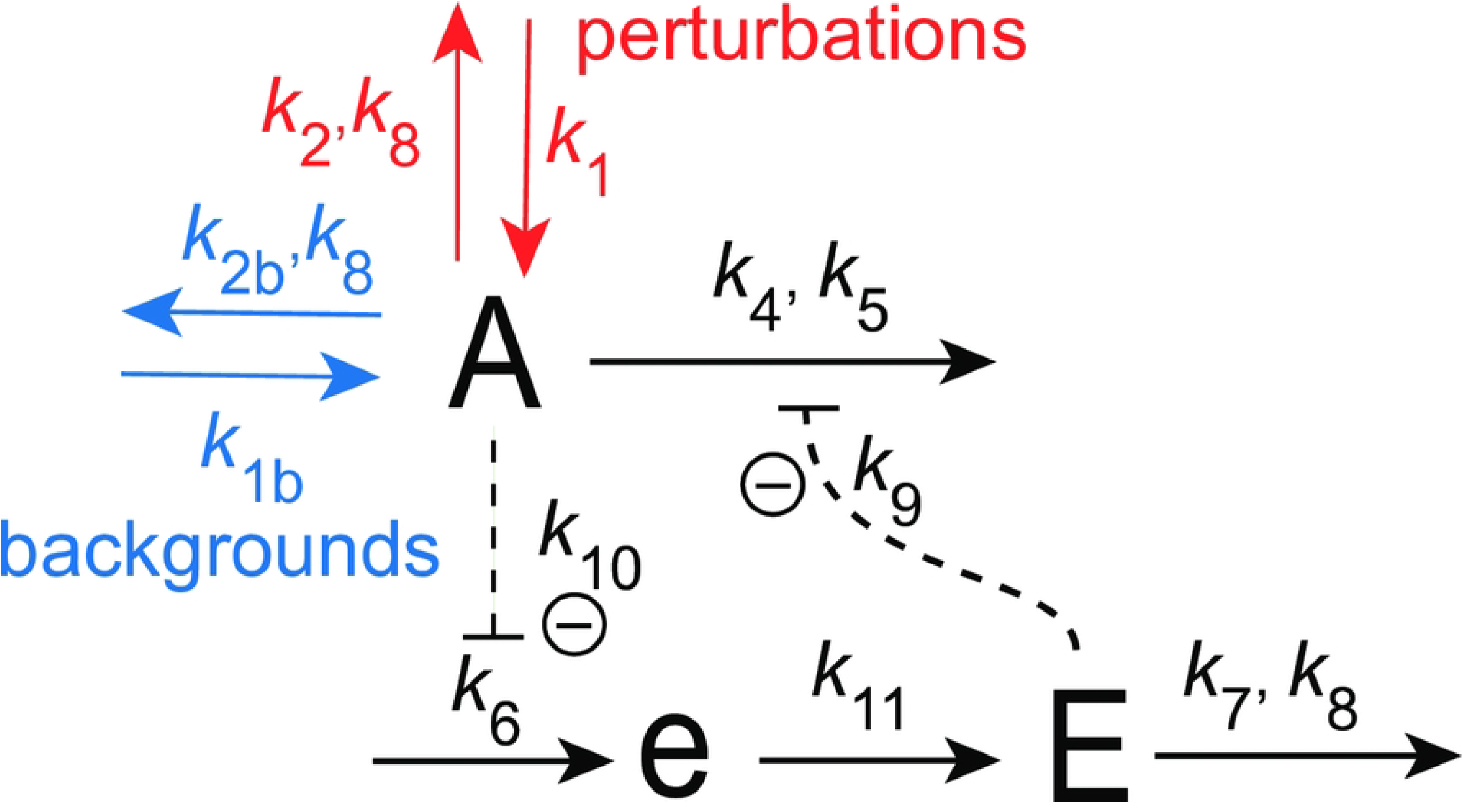
Motif 8 based negative feedback loop with intermediate *e*. Reactions outlined in red undergo perturbative steps or pulses, while reactions indicated in blue represent constant backgrounds.

This controller is also based on derepression by the manipulated variable *E*, but acts as an outflow controller, i.e. compensates for inflow perturbations to *A*. Unlike the M2 controller in Fig 2 this feedback loop does not control the concentration of *A*, but keeps the *A*-regulated flux to *e* constant. The rate equations are:

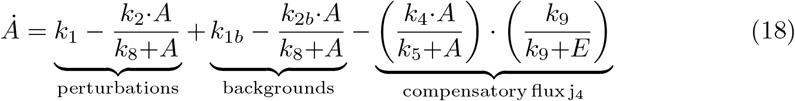

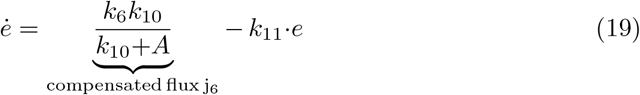

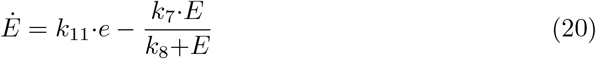

Due to zero-order removal of *E* (*k*_8_ ≪ *E*) the average flux *<j*_6_*>* = *<k*_6_*k*_10_*/*(*k*_10_+*A*)*>* is under homeostatic control. This is seen by setting Eqs 19 and 20 to zero and eliminating the term *k*_11_·*e*. This leads to:

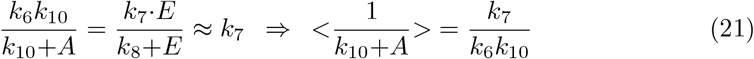

Thus, instead of controlling *<A>* the feedback in Fig 16 controls the property *<*1*/*(*k*_10_+*A*)*>*, which is proportional to the average flux *<j*_6_*>*.

Apart from the condition by Ang et al. [44] referred above, the controller has two operational limits:

i. a capacity limit to compensate inflows to *A* and
ii. the controller’s inability to compensate dominating outflows from *A*.

When *k*_2_=*k*_2*b*_=0, the maximum allowable inflow to *A* is *k*_1_+*k*_1*b*_, which is balanced by the maximum possible compensatory flux *j*_4_=*k*_4_ when *E* → 0, i.e. when *j*_4_ → *k*_4_. As the total outflow *k*_2_+*k*_2*b*_ increases the total inflow to *A, k*_1_+*k*_1*b*_, can increase accordingly. However, an outflow controller is not able to compensate outflow perturbations, which implies that controller breakdown will occur whenever *k*_2_+*k*_2*b*_ ≥ *k*_1_+*k*_1*b*_.

These two scenarios of controller breakdown are presented in Fig 17. In Fig 17a the concentrations of *A* (left panel) and *E* (right panel) are shown when *k*_1_ increases successively during three phases starting with *k*_1_=5 × 10^2^ (phase 1), to *k*_1_=8 × 10^3^ (phase 2), and finally in phase 3 to *k*_1_=1.5 × 10^4^ while *k*_2_=*k*_1*b*_=*k*_2*b*_=0.0. In phase 3 *k*_1_ exceeds the maximum compensatory flux of *k*_4_=1.0 × 10^4^ and the controller breaks down: concentration *A* increases rapidly while *E* decreases. The left panel in Fig 17a shows in addition, outlined in blue, the calculated value of *<*1*/*(*k*_10_+*A*)*>*. Its setpoint, *k*_7_*/*(*k*_6_*k*_10_)=0.5, is indicated by the thick orange line.

**Fig 17.**
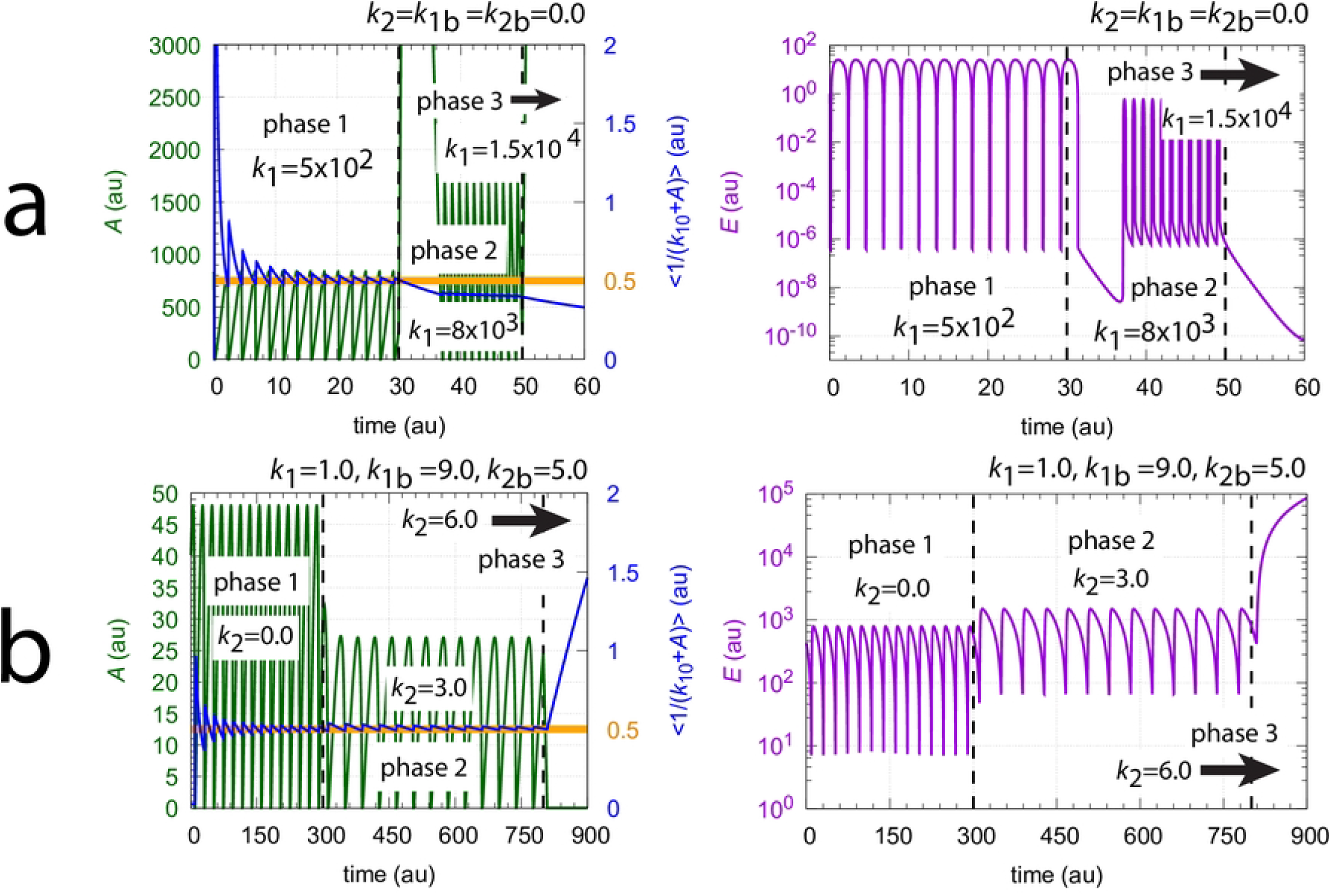
Breakdown of controller Fig 16 by (a) exceeding the capacity of the compensatory flux *j*_4_, and (b) by the dominance of the outflow rate from *A* with respect to inflows to *A*. Rate constants, panel a: *k*_1_ (phase 1)=5×10^2^, *k*_1_ (phase 2)=8×10^3^, *k*_1_ (phase 3)=1.5×10^4^, *k*_1*b*_=*k*_2_=*k*_2*b*_=0.0, *k*_4_=1.0×10^4^, *k*_5_=1.0×10^−6^, *k*_6_=1.0×10^3^, *k*_7_=50.0, *k*_8_=1.0×10^−6^, *k*_9_=*k*_10_=0.1, *k*_11_=1.0. Initial concentrations, panel a: *A*_0_=837.94, *E*_0_=1.3246, *e*_0_=15.514. Rate constants, panel b: *k*_1_=1.0, *k*_1*b*_=9.0, *k*_2_ (phase 1)=0.0, *k*_2_ (phase 2)=3.0, *k*_2_ (phase 3)=6.0, *k*_2*b*_=5.0. Other rate constants as in figure a. Initial concentrations, panel b: *A*_0_=39.942, *E*_0_=430.14, *e*_0_=2.7212.

Fig 17b shows the controller’s breakdown when the total outflow from *A* becomes dominant. Here we have constant levels of *k*_1_=1.0, *k*_1*b*_=9.0, and *k*_2*b*_=5.0. As *k*_2_ increases from 0.0 (phase 1) to 3.0 (phase 2), and finally in phase 3 to 5.0, the controller breaks down in the attempt to force the compensatory flux *j*_4_ to zero by a steady increase (windup) of *E*. The blue line in the left panel of Fig 17b shows the homeostasis in *<*1*/*(*k*_10_+*A*)*>* during phases 1 and 2 and its breakdown in phase 3.

#### Phase response curves of single-loop feedback M8

Next I show how the PRCs of the single-loop feedback in Fig 16 behave as backgrounds change. Figs 18 and 19 show two sets of calculations where *k*_1*b*_ and *k*_2*b*_ are changed,respectively. The final phase shifts are outlined in orange. Together with the PRC, one cycle of the undisturbed oscillation is shown (outlined in blue) to see the PRC in relationship with the oscillation and the period length. Interestingly, the single-loop M8 oscillator appears to have only positive phase shifts. A direct comparison between the PRCs is made in Fig 20 using a normalized phase of perturbation. The maximum PRC amplitude decreases with increasing *k*_1*b*_ background (Fig 20a), which is reminiscent of Weber’s law which was observed earlier for this controller (see Fig 11 in [54]). In other words: the response amplitude is diminished at increased backgrounds which are applied in parallel to a constant perturbation to which the controller opposes to. On the other hand, the increase of the PRC amplitude at increased *k*_2*b*_’s represents the ‘opposite’ situation: at increased *k*_2*b*_ the controller needs to reduce less *E* concentrations (or less *<E>*) in order to oppose identical perturbations in *k*_1_.

**Fig 18.**
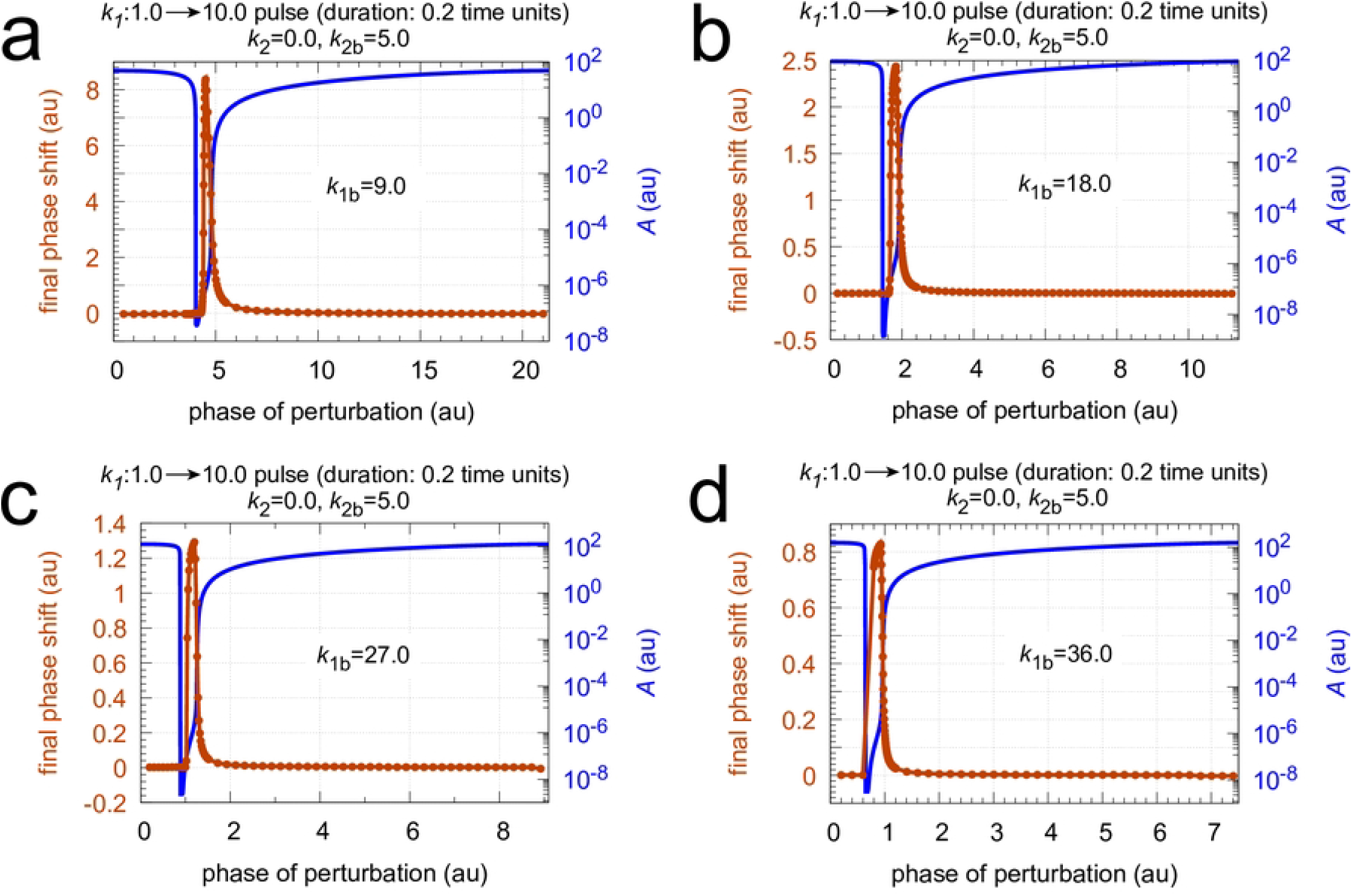
Phase response curves of oscillator Fig 16 for *k*_1_ pulses 1.0 → 10.0 with a duration of 0.2 time units at different *k*_1*b*_ backgrounds. Panel a: *k*_1*b*_=9.0; panel b: *k*_1*b*_=18.0; panel c: *k*_1*b*_=27.0; panel d: *k*_1*b*_=36.0. Note the successive decrease in the maximum phase response amplitude as *k*_1*b*_ increases. Rate constants *k*_2_=0.0 and *k*_2*b*_=5.0. Other rate constants as in Fig 17. Initial concentrations: panel a, *A*_0_=48.056, *E*_0_=199.85, *e*_0_=2.1163; unperturbed period=21.3. Panel b, *A*_0_=92.430, *E*_0_=71.292, *e*_0_=1.1734; unperturbed period=11.4. Panel c, *A*_0_=126.32, *E*_0_=43.311, *e*_0_=1.0321; unperturbed period=9.0. Panel d, *A*_0_=155.52, *E*_0_=31.108, *e*_0_=1.1462; unperturbed period=7.5. All initial concentrations start at an *A* maximum.

**Fig 19.**
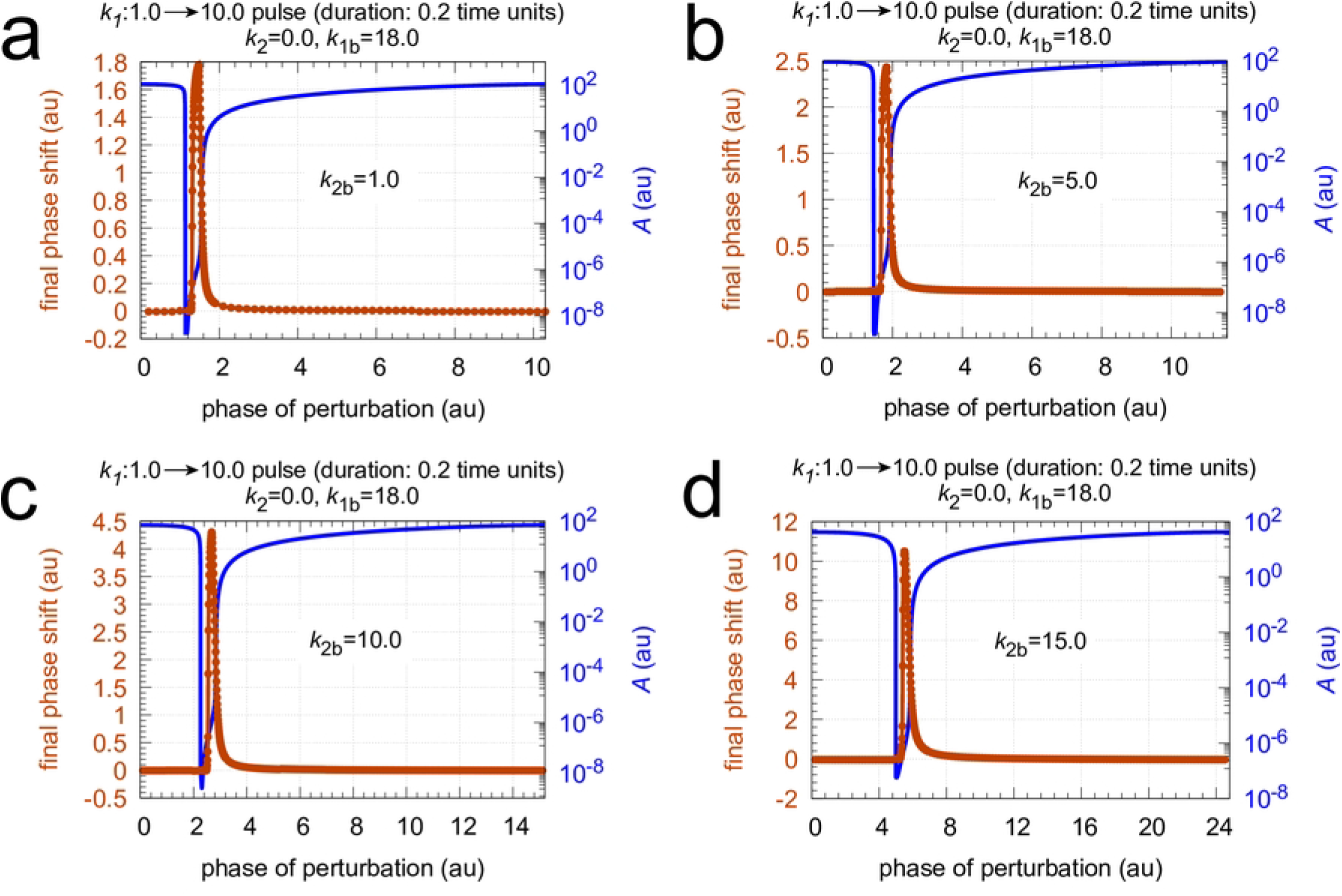
Phase response curves of oscillator in Fig 16 for *k*_1_ pulses 1.0 → 10.0 with a duration of 0.2 time units at different *k*_2*b*_ backgrounds. Panel a: *k*_2*b*_=1.0; panel b: *k*_2*b*_=5.0; panel c: *k*_2*b*_=10.0; panel d: *k*_2*b*_=15.0. Note the now successive increase in the maximum phase response amplitude as *k*_2*b*_ increases. Rate constants *k*_2_=0.0 and *k*_1*b*_=18.0. Other rate constants as in Fig 17. Initial concentrations: panel a, *A*_0_=108.26, *E*_0_=55.385, *e*_0_=1.0664; unperturbed period=10.6. Panel b, *A*_0_=92.430, *E*_0_=71.322, *e*_0_=1.1734; unperturbed period=11.5. Panel c, *A*_0_=70.007, *E*_0_=110.01, *e*_0_=1.4837; unperturbed period=15.2. Panel d, *A*_0_=41.687, *E*_0_=249.90, *e*_0_=2.4280; unperturbed period=24.7. All initial concentrations start at an *A* maximum.

**Fig 20.**
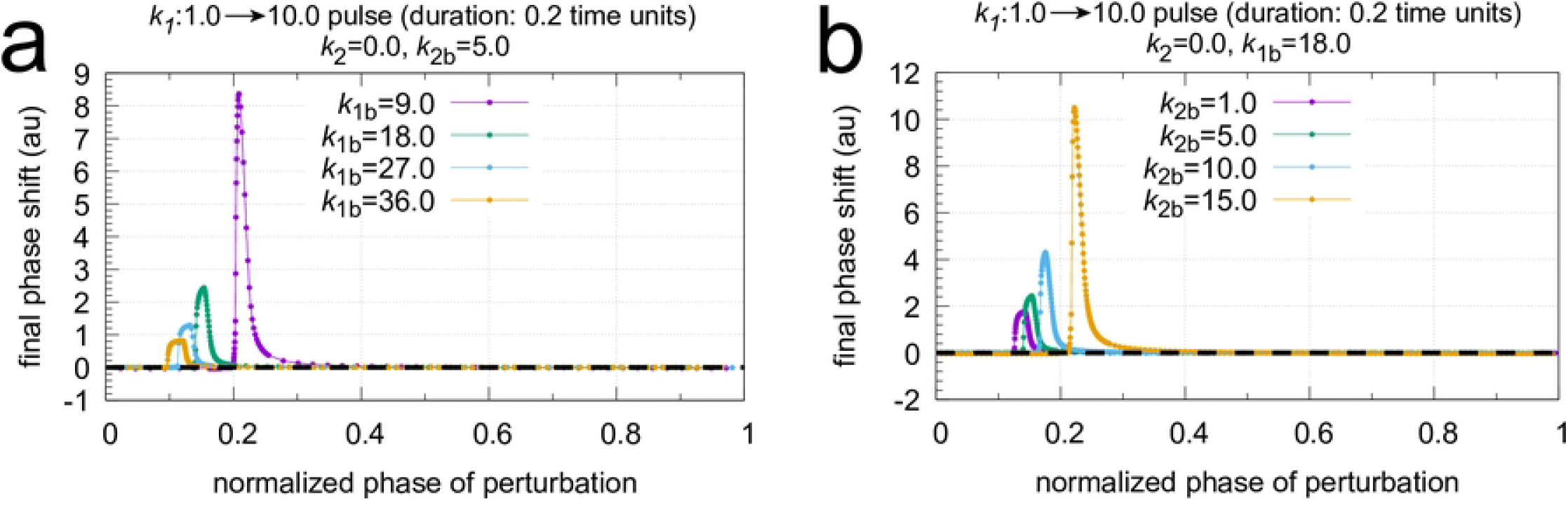
Influence of *k*_1*b*_ and *k*_2*b*_ backgrounds on the phase response curves of the oscillator in Fig 16. To make a direct comparison possible the phases of stimulation are normalized with respect to the oscillator’s period length (see Figs 18 and 19). Panel a: comparing phase response curves of Fig 18. Panel b: comparing phase response curves of Fig 19.

#### M8 coherent feedback

Fig 21 shows the considered coherent feedback scheme with the M8 feedback from Fig 16 in the center. As in Fig 4 we have that *I*_1_ and *I*_2_ act as regulators to keep also *<E>* under homeostatic control, which makes the frequency of the oscillator robust against perturbations [29].

**Fig 21.**
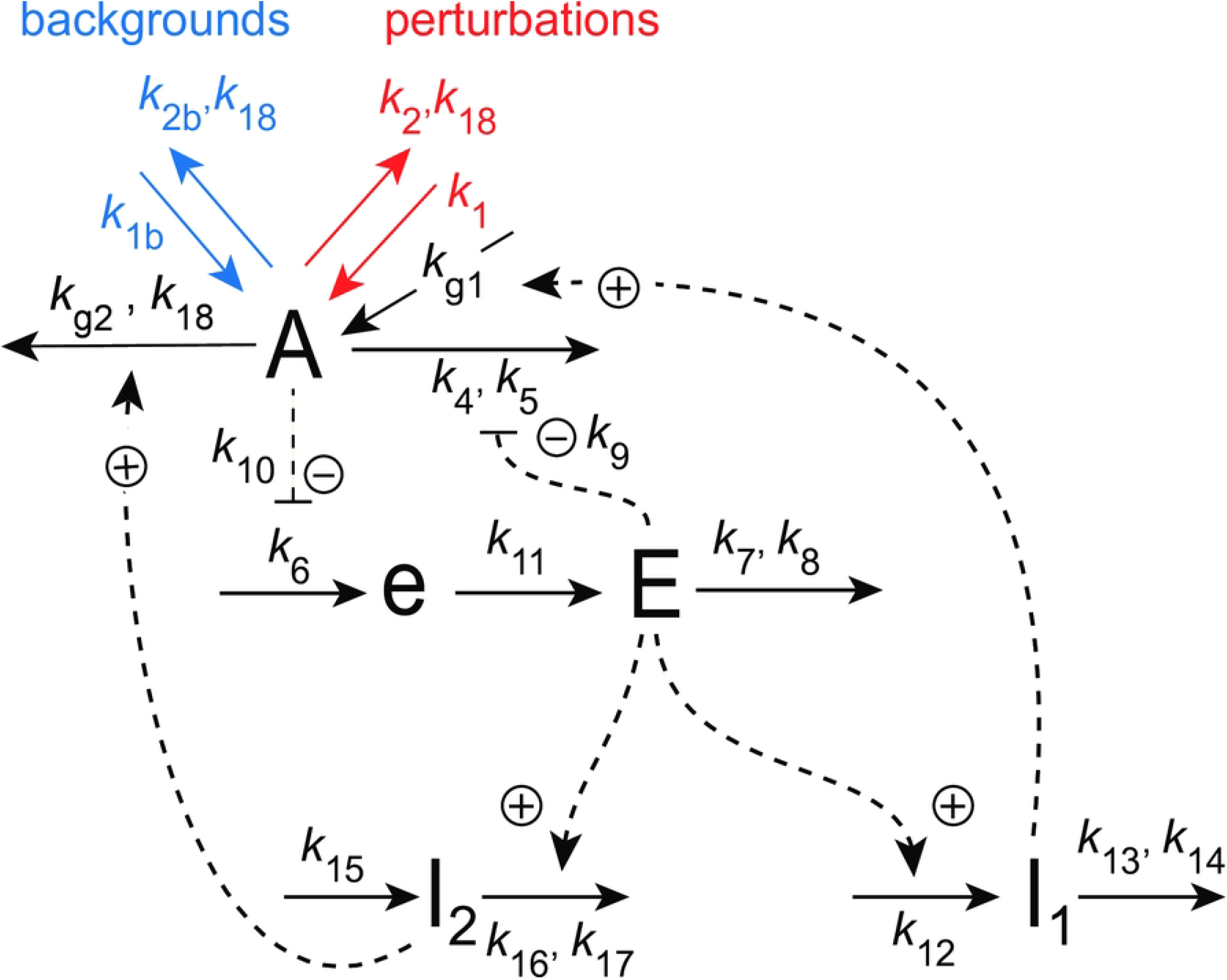
Coherent feedback scheme using *I*_1_ and *I*_2_, which control the level of *<j*_6_*>* and *<E>* in the central M8-type feedback loop (Fig 16).

The rate equations are:

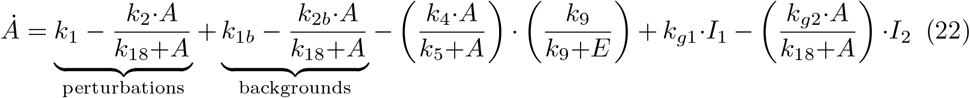

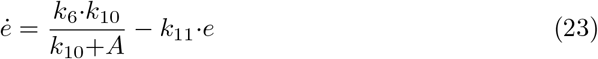

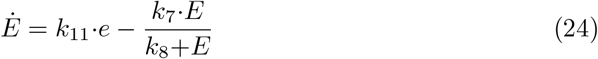

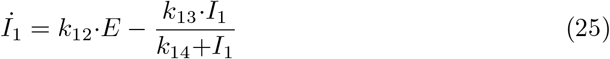

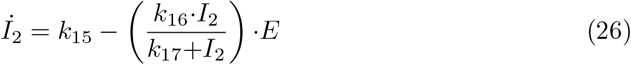

As for the coherent M2 oscillator in Fig 4 we have two setpoints for *<E>*. From the steady state condition of Eqs 25 and 26 together with the zero-order removals of *I*_1_ and *I*_2_ (i.e. *I*_1_*/*(*k*_14_+*I*_1_)≈1 and *I*_2_*/*(*k*_17_+*I*_2_)≈1) we get:

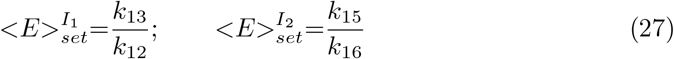

Eliminating *k*_11_·*e* from the steady state expressions of Eqs 23 and 24 leads to the flux control of *j*_6_=*k*_6_*k*_10_*/*(*k*_10_+*A*) by

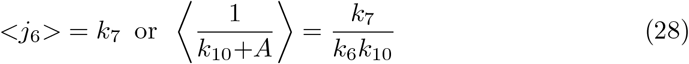

Fig 22 shows the behavior of the M8 coherent feedback when the same perturbation and background conditions are applied as in Fig 17b. The breakdown of the controller at *k*_2_=6.0 as seen there is now avoided, because *I*_1_ is able to compensate for the excess outflow from *A* (panel c). Panel a, right ordinate shows the homeostasis in *<*1*/*(*k*_10_+*A*)*>*, while panels b and d show homeostasis in *<E>* and the frequency, respectively. Even *<A>* appears to be under homeostatic control, although there is no explicit mathematical expression for its setpoint!

**Fig 22.**
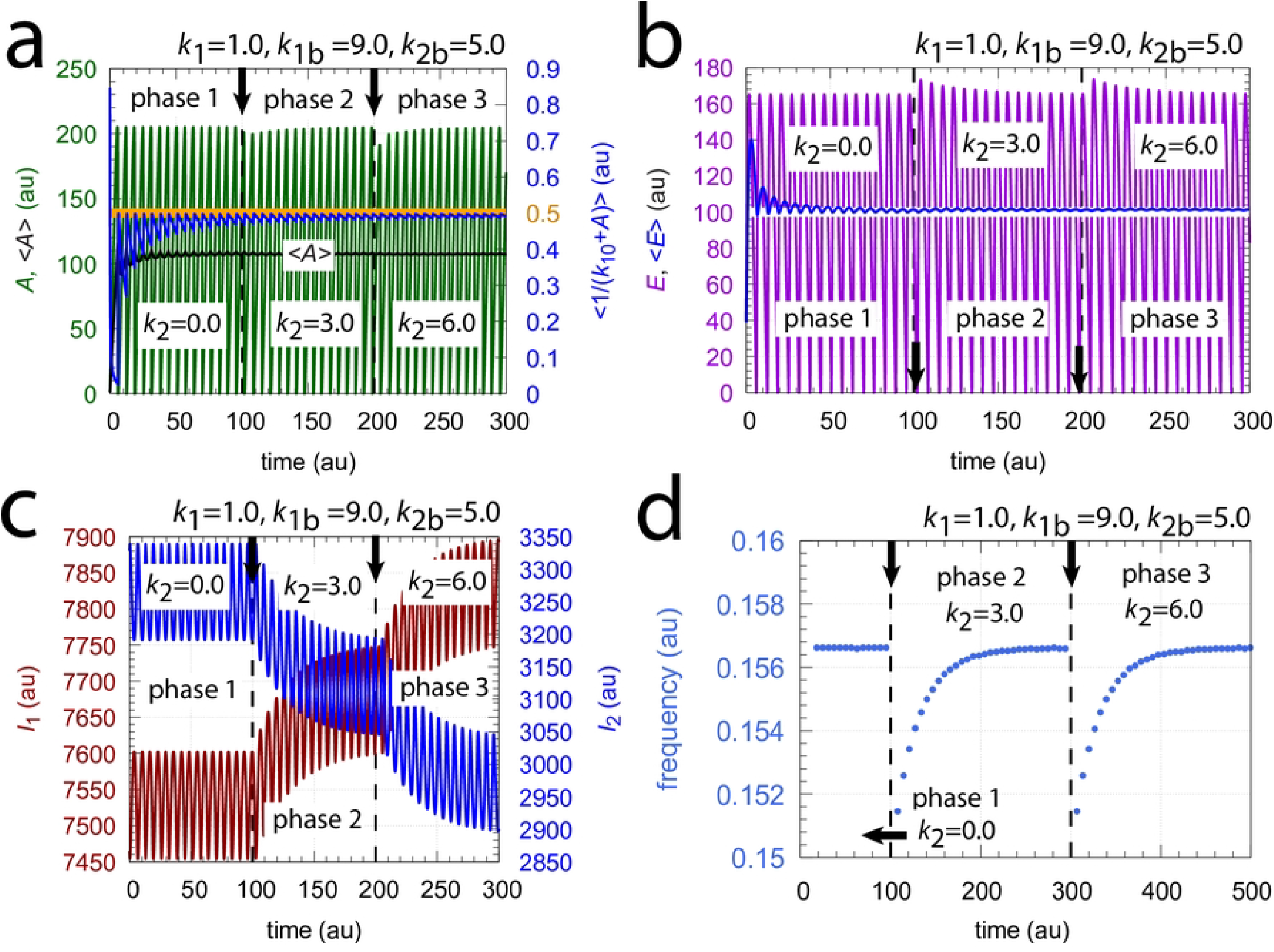
Homeostasis in frequency, in the average flux *<*1*/*(*k*_10_+*A*)*>*, and in the average of *<E>* by coherent feedback scheme of Fig 21. At times *t*=100 and *t*=200, indicated by the vertical arrows, *k*_2_ : 0.0 → 3.0 and *k*_2_ : 3.0 → 6.0 steps were applied, respectively with *k*_1_=1.0 and backgrounds *k*_1*b*_=9.0 & *k*_2*b*_=5.0. Panel a, left ordinate: *A* (outlined in green) and *<A>* (outlined in black) as functions of time. Panel a, right ordinate: flux *<*1*/*(*k*_10_+*A*)*>* (outlined in blue) as a function of time. Orange line and number indicate the setpoint *<*1*/*(*k*_10_+*A*)*>*_*set*_=*k*_7_*/*(*k*_6_*k*_10_)=0.5. Panel b: *E* (outline in purple) and *<E>* (outlined in blue) as a function of time. The white line indicates the setpoint *<E>*_*set*_=*k*_15_*/k*_16_=*k*_13_*/k*_12_=100.0. Panel c: *I*_1_ and *I*_2_ (outlined respectively in red and blue) as a function of time. Panel c: frequency as a function of time. Other rate constants: *k*_4_=1.0 × 10^4^, *k*_5_=1.0 × 10^−6^, *k*_6_=1.0 × 10^3^, *k*_7_=50.0, *k*_8_=1.0 × 10^−6^, *k*_9_=*k*_10_=0.1, *k*_11_=1.0, *k*_12_=1.0, *k*_13_=100.0, *k*_14_=1.0 × 10^−6^, *k*_15_=100.0, *k*_16_=1.0, *k*_17_=*k*_18_=1.0 × 10^−6^, *k*_*g*1_=*k*_*g*2_=0.01. Initial concentrations: *A*_0_=0.9913, *E*_0_=37.985, *e*_0_=247.94, *I*_1,0_=7.466 × 10^3^, and *I*_2,0_=3.328 × 10^3^. All averages were calculated by Eq 2.

The coherent M8 feedback scheme shows background compensation when the frequency resetting is tested. Fig 23 shows two resettings, one for *k*_1_ : 1.0 → 10.0 steps (panel a) and the other (panel b) for *k*_2_ : 0.0 → 10.0 steps. For both perturbation types the large orange dots relate to the background combination *k*_1*b*_=90.0, *k*_2*b*_=5.0 while for the smaller blue dots have the backgrounds *k*_1*b*_=9.0, *k*_2*b*_=50.0. The panels to the right show the projections of all frequency-time data on to the frequency-time plane. Background compensation is indicated by the complete alignment of the two background combinations.

**Fig 23.**
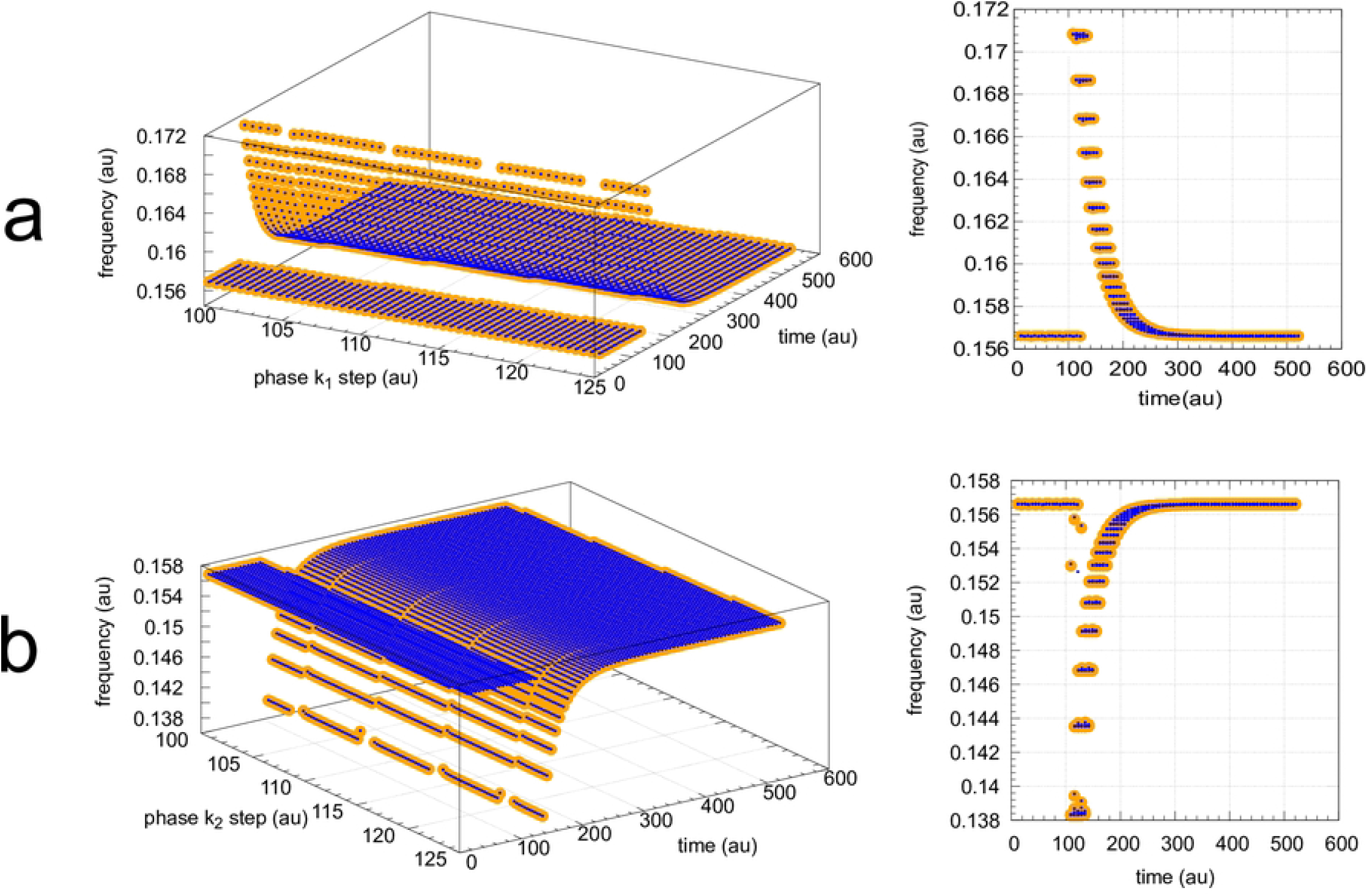
Background compensation in the frequency resetting of the m8-type oscillator Fig 21. Figure a, left panel: frequency resetting for *k*_1_ : 1.0 → 10.0 steps applied at times *t*=100.0 until *t*=125.0 by steps of 0.5 time units. Backgrounds: large orange dots, *k*_1*b*_=90.0 & *k*_2*b*_=5.0; small blue dots, *k*_1*b*_=9.0 & *k*_2*b*_=50.0. Figure a, right panel: same data as in left panel, but showing the projections on the frequency-time plane. Initial concentrations (starting at a maximum of *A*): *A*_0_=204.88, *E*_0_=20.763, *e*_0_=1.563, *I*_1,0_=3.485×10^3^, *I*_2,0_=7.308×10^3^. *k*_2_=1.0, other rate constants as in Fig 22. Figure b, left panel: frequency resetting for *k*_2_ : 0.0 → 10.0 steps applied at times *t*=100.0 until *t*=125.0 by steps of 0.25 time units. Figure b, right panel: same data as in left panel, but showing the projections on the frequency-time plane. Initial concentrations as in figure a. *k*_1_=1.0, other rate constants as in Fig 22.

Finally, I looked at the phase shifts for the *k*_1_ and *k*_2_ perturbations and background combinations given in Fig 23. An example how final phase shifts have been determined is illustrated in Fig 24. In panel a, a *k*_1_ : 1.0 → 10.0 step is applied at phase *t*=3.0 showing the response for the two backgrounds *k*_1*b*_=90.0 & *k*_2*b*_=5.0 and *k*_1*b*_=9.0 & *k*_2*b*_=50.0. In panel b a *k*_2_ : 0.0 → 10.0 step is applied at the same phase as in panel a testing the same two backgrounds. As for the frequency in Fig 23 the transient and final phase shifts ΔΦ are independent of the two backgrounds. In fact, the phase response curves (Fig 25) show constant final phase shifts which are background compensated.

**Fig 24.**
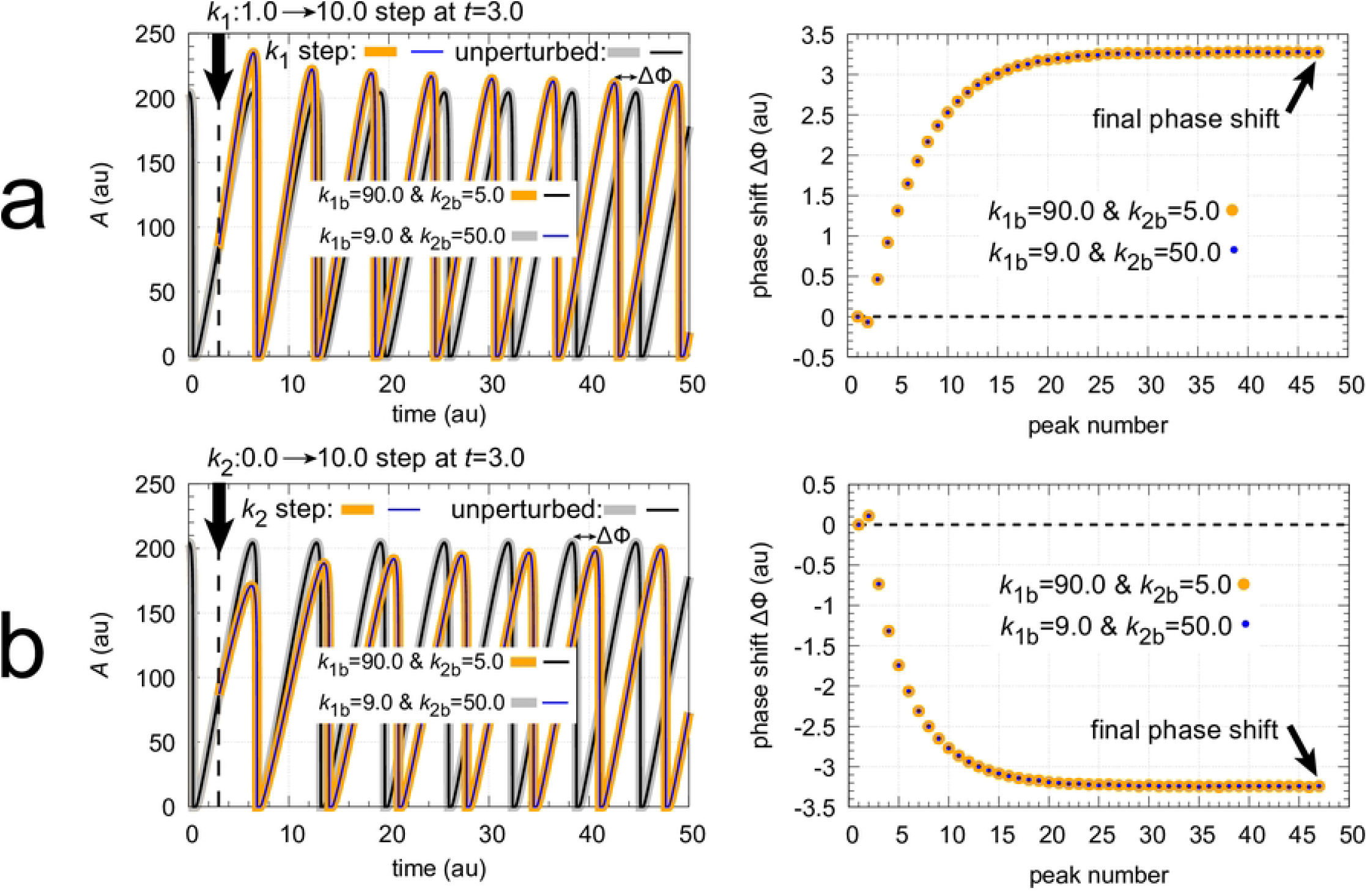
Background independence of phase shifts in the coherent feedback scheme Fig 21 towards *k*_1_ and *k*_2_ steps. Figure a, left panel: Concentration of *A* at the two background combinations (*k*_1*b*_=90.0 & *k*_2*b*_=5.0) and (*k*_1*b*_=9.0 & *k*_2*b*_=50.0) when a *k*_1_ : 1.0 → 10.0 step indicated by the vertical arrow is applied at *t*=3.0. For the background combination *k*_1*b*_=90.0 & *k*_2*b*_=5.0 the thin black line shows *A* of the unperturbed oscillator, while the thick orange line shows the effect of the *k*_1_ step. Initial concentrations are: *A*_0_=204.34, *E*_0_=20.838, *e*_0_=1.566, *I*_1,0_=4.935×10^4^, *I*_2,0_=5.319×10^4^. For the background combination *k*_1*b*_=9.0 & *k*_2*b*_=50.0, the thick gray line shows the unperturbed oscillator, and the thin blue line shows the effect of the *k*_1_ step. Initial concentrations are: *A*_0_=204.36, *E*_0_=20.846, *e*_0_=1.567, *I*_1,0_=5.565×10^4^, *I*_2,0_=4.689×10^4^. The phase shift Δ Φ between unperturbed and perturbed peaks is indicated in the upper right corner of the graph. Rate constant *k*_2_=0.0. The right panel of figure a shows that the transient phase shifts shown as a function of peak number are independent of the two backgrounds. Figure b, left panel: Concentration of *A* for the two background combinations above, but a *k*_2_ : 0 → 10.0 step is now applied at *t*=3.0. Initial concentrations as for figure a. Rate constant *k*_1_=1.0. The right panel of figure b shows that the phase shifts are independent of the two background combinations. Other rate constants as in Fig 22.

**Fig 25.**
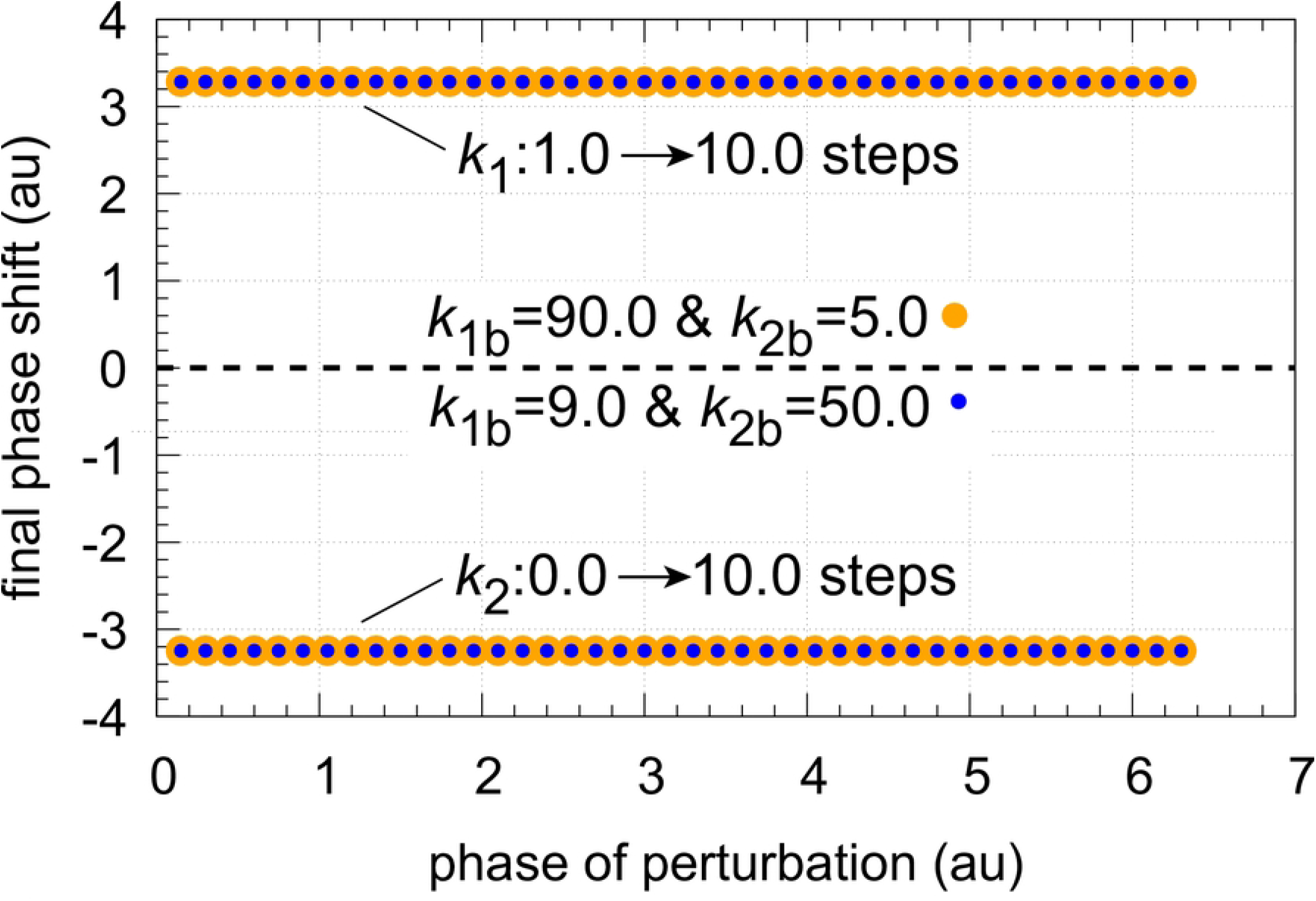
Final phase shift (see definition Fig 24, right panels) as a function of phase of perturbation for *k*_1_ : 1.0 → 10.0 and *k*_2_ : 0.0 → 10.0 steps at backgrounds *k*_1*b*_ = 90.0 & *k*_2*b*_ = 5.0 (orange dots) and *k*_1*b*_ = 9.0 & *k*_2*b*_ = 50.0 (blue dots). Initial concentrations and rate constants as in Fig 24.

### Summary and outlook

I have shown that coherent feedback oscillators have the ability to compensate their frequency resetting and phase shifts against different but constant backgrounds. This indicates that these systems, either oscillatory or nonoscillatory [18] seem to have the potential to ‘ignore’ ambient noise. Classical mechanisms to deal with ambient noise can be an increase of the call amplitude, known as the Lombard Effect [56, 57], or a change of the call frequency [58]. However, the mechanism of background compensation by coherent feedback is different, as it actively compensates for the background. To what extent background compensation considered here is being used in organisms which live in noisy environments is not known, but it appears interesting to do research in this direction.

## Supporting information

**S1 Programs. Python scripts**. A zip-file with Python and Matlab scripts showing results from Figs 3, 5, 7, 8, 12, 14, 17, 22, and 24.

**S1 Movies. Zip file containing animations of Fig 9**. The Quicktime movie files show the trajectories in phase space when *k*_1_ and *k*_2_ step perturbations are applied.

The moving cursor has a length of 1 time unit. The movies also show the preservations of the limit cycles after the step perturbations when limit cycles are projected on to the *A*-*E* phase space.

**S2 Movie. Animation of Fig 10**. The movie shows Fig 10 from various viewing angles.

**S3 Movie. Animation of Fig 11**. The movie shows Fig 11 from various viewing angles.

## Notes

### Competing Interest Statement

The authors have declared no competing interest.

